# Scalable pooled CRISPR screens with single-cell chromatin accessibility profiling

**DOI:** 10.1101/2020.11.20.390971

**Authors:** Noa Liscovitch-Brauer, Antonino Montalbano, Jiale Deng, Alejandro Méndez-Mancilla, Hans-Hermann Wessels, Nicholas G. Moss, Chia-Yu Kung, Akash Sookdeo, Xinyi Guo, Evan Geller, Suma Jaini, Peter Smibert, Neville E. Sanjana

## Abstract

Pooled CRISPR screens have been used to identify genes responsible for specific phenotypes and diseases, and, more recently, to connect genetic perturbations with multi-dimensional gene expression profiles. Here, we describe a method to link genome-wide chromatin accessibility to genetic perturbations in single cells. This scalable, cost-effective method combines pooled CRISPR perturbations with a single-cell combinatorial indexing assay for transposase-accessible chromatin (CRISPR-sciATAC). Using a human and mouse species-mixing experiment, we show that CRISPR-sciATAC separates single cells with a low doublet rate. Then, in human myelogenous leukemia cells, we apply CRISPR-sciATAC to target 21 chromatin-related genes that are frequently mutated in cancer and 84 subunits and cofactors of chromatin remodeling complexes, generating chromatin accessibility data for ~30,000 single cells. Using this large-scale atlas, we correlate loss of specific chromatin remodelers with changes in accessibility — globally and at the binding sites of individual transcription factors. For example, we show that loss of the H3K27 methyltransferase EZH2 leads to increased accessibility at heterochromatic regions involved in embryonic development and triggers expression of multiple genes in the *HOXA* and *HOXD* clusters. At a subset of regulatory sites, we also analyze dynamic changes in nucleosome spacing upon loss of chromatin remodelers. CRISPR-sciATAC is a high-throughput, low-cost single-cell method that can be applied broadly to study the role of genetic perturbations on chromatin in normal and disease states.

---

Chromatin accessibility orchestrates *cis*- and *trans*-regulatory interactions to control gene expression and is dynamically regulated in cell differentiation and homeostasis. Alterations in chromatin state have been associated with many diseases including several cancers^1^. To study how genetic perturbations affect chromatin states, we developed a novel platform for scalable pooled CRISPR screens with single-cell ATAC-seq profiles: CRISPR-sciATAC. In CRISPR-sciATAC, we simultaneously capture Cas9 single-guide RNAs (sgRNAs) and perform single-cell combinatorial indexing ATAC-seq^2^ (**Fig. 1a**, **Supplementary Fig. 1**). Following cell fixation and lysis, nuclei are recovered and the open chromatin regions of the genomic DNA undergo barcoded tagmentation in a 96-well plate using a unique, easy-to purify transposase from *Vibrio parahemolyticus* (**Fig. 1b**, **Supplementary Fig. 2**). Next, the sgRNA is barcoded with the same barcode sequence as the ATAC fragments, using *in situ* reverse transcription. The nuclei are pooled together and split again to a new 96-well plate and both the ATAC fragments and the sgRNA are tagged with a second barcode in two consecutive PCR steps. At the end of this process, a unique combination of barcodes (“cell barcode”) tag both the sgRNA and the ATAC fragments from each cell (**Fig. 1a**, **Supplementary Fig. 1, Supplementary Table 1**).

**Fig. 1.**
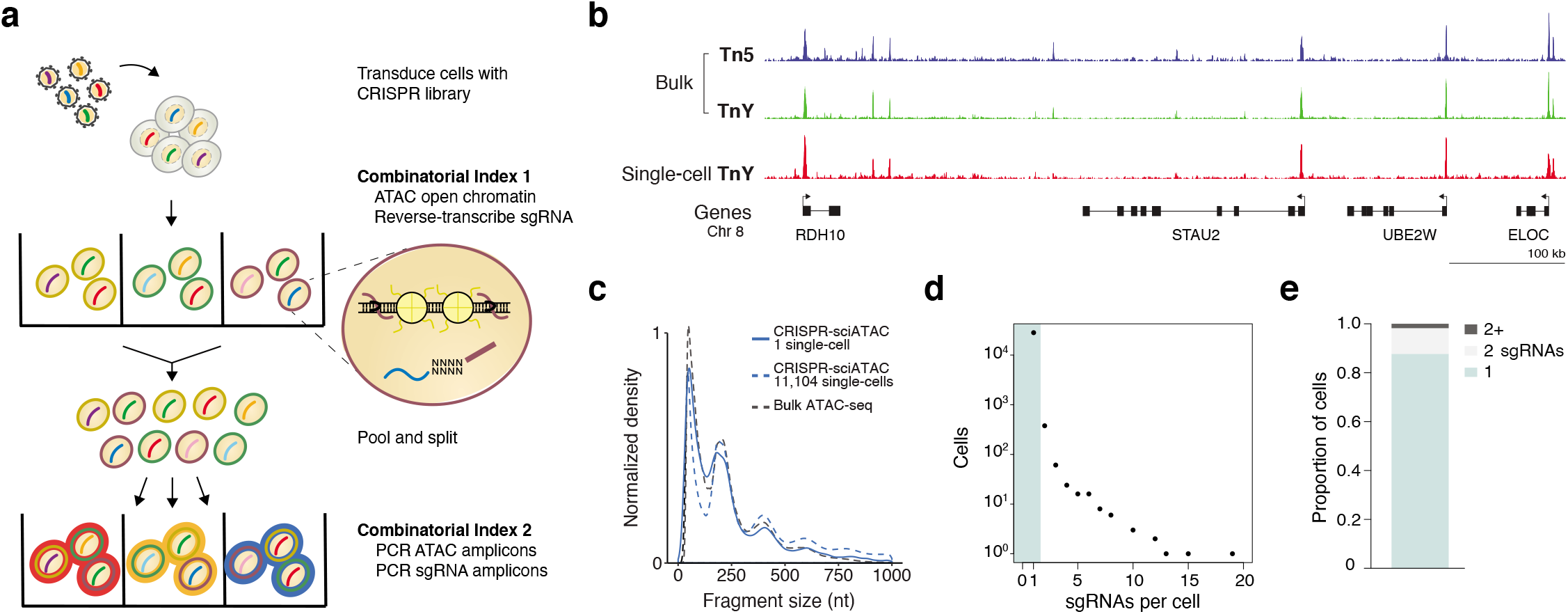
CRISPR screens with single-cell combinatorial indexing assay of transposable and accessible chromatin sequencing (CRISPR-sciATAC) enables the joint capture of chromatin accessibility profiles and CRISPR perturbations. (**a**) CRISPR-sciATAC workflow with initial barcoding, nuclei pooling and re-splitting, and then second round barcoding. (**b**) Comparison of bulk ATAC-seq chromatin accessibility profiles from K562 cells using Tn5 and TnY transposases and aggregated CRISPR-sciATAC single cell profiles from 11,104 cells. (**c**) ATAC-seq fragment size distribution from K562 cells of bulk ATAC-seq data, aggregated CRISPR-sciATAC single cell profiles from 11,104 cells and one representative single cell from CRISPR-sciATAC. (**d**) Number of CRISPR single-guide RNAs (sgRNAs) detected per cell. (**e**) Proportion of cells with 1, 2, or more than 2 sgRNAs.

To quantify capture and barcoding of single cells, we performed CRISPR-sciATAC on a mix of human (HEK293) and mouse (NIH3T3) cells. Human and mouse cells were each transduced with a small library of 10 distinct non-targeting sgRNAs with no overlapping sgRNAs between the two pools. The guide sequences were cloned into a lentiviral vector that includes the guide RNA within a Pol2 transcript (CROP-seq vector)^3^. We found that 93% of cell barcodes had sgRNA-containing reads that could uniquely be assigned to either human or mouse sgRNAs (**Supplementary Fig. 3a**) and 96% of cell barcodes had ATAC-seq reads mapping to either the human or mouse genome, indicating that the majority of cell barcodes were correctly assigned to single cells (**Supplementary Fig. 3b**). As an additional verification of single-cell separation, we also measured the species concordance between the ATAC-seq and sgRNA reads. We found that for 92% of the captured cell barcodes both ATAC-seq and sgRNA reads aligned either to human or mouse reference genomic and sgRNA sequences, respectively. In 4.4% of cells, the ATAC-seq and/or sgRNA reads could not be exclusively assigned to one species. ATAC-seq and sgRNA reads were assigned to different species (species collision) in 3.6% of cells (**Supplementary Fig. 3c**). The low rates of these two failure modes suggest that CRISPR-sciATAC can simultaneously identify accessible chromatin and CRISPR sgRNAs in single cells.

To test the ability of CRISPR-sciATAC to capture biologically meaningful changes in chromatin accessibility, we targeted 21 chromatin modifiers that are highly mutated in cancer (**Supplementary Fig. 4a, b**). Using the Catalog of Somatic Mutations in Cancer (COSMIC) database^4^, we selected 21 chromatin-related genes that carry the highest mutational load across all cancers, including 9 chromatin remodelers (*ARID1A, ATRX, CHD4, CHD5, CHD8, MBD1, PBRM1, SMARCA4*, and *SMARCB1*), 2 DNA methyltransferases (*DNMT3A* and *TET2*), 3 histone methyltransferases (*EZH2, PRDM9*, and *SETD2*), 1 histone demethylase (*KDM6A*), 1 histone deacetylase (*HDAC9*), 3 histone subunits (*H3F3A, H3F3B*, and *HIST1H3B*) and 2 readers (*ING1* and *PHF6*). We designed 3 sgRNAs per gene and also included 3 non-targeting sgRNAs in our library (**Supplementary Table 2**). We transduced Cas9-expressing human myelogenous leukemia K562 cells with this lentiviral sgRNA library at a low multiplicity of infection and selected with puromycin for transduced cells. After 1 week of selection, we collected single-cell paired-end ATAC-seq data. After filtering for cells with ≥500 unique ATAC-seq fragments and ≥100 sgRNA reads (**Supplementary Fig. 4c-f**), we obtained 11,104 cells with a median of 1,977 unique ATAC-seq fragments mapping to the human genome. Aggregated ATAC-seq profiles for these cells correlate well with bulk data from K562 cells (**Fig. 1b**, **Supplementary Fig. 4g**). Single cells retained an ATAC fragment length distribution similar to cells tagmented in bulk (**Fig. 1c**). The majority of cell barcodes (83%) had one sgRNA (**Fig. 1d, e**), and for 90% of cell barcodes, a single sgRNA represented ≥99% of the reads (**Supplementary Fig. 4h**).

We recovered all 66 sgRNAs with a median of 148 single cells per sgRNA and 468 single cells per gene. Upon closer examination, we noticed that not all gene targets resulted in the same number of single cells captured, suggesting that some of our targets might be essential genes whose targeting leads to drop-out of those cells. To distinguish sgRNA depletion of essential genes from inability to capture sgRNAs using CRISPR-sciATAC, we separately amplified sgRNAs from the bulk population at an early time point and at 1 and 2 weeks post-selection (**Supplementary Fig. 5a**). We found high correlation between all samples across 3 independent transduction replicates (**Supplementary Fig. 5b, c**). For several genes, multiple, distinct sgRNAs targeting the same gene were consistently depleted or enriched: *H3F3A, CHD4, SMARCA4*, and *SMARCB1* were depleted, while targeting *KDM6A* accelerated cell growth (**Supplementary Fig. 5d**). Using robust rank aggregation to measure consistent enrichment across multiple sgRNAs^5^, we computed gene-level enrichment scores (**Supplementary Fig. 5e, Supplementary Table 3**), which were highly correlated with a previous genome-wide CRISPR screen in K562 cells^6^ (*r* = 0.85) (**Supplementary Fig. 5f**). Reassuringly, enrichment of individual sgRNAs was positively correlated with cell numbers estimated from CRISPR-sciATAC cell barcodes (*r* = 0.73, Supplementary **Fig. 5g**). Different sgRNAs targeting the same gene tend to result in similar numbers of single cells, highlighting consistent proliferation phenotypes between different genetic perturbations targeting the same gene (**Supplementary Fig. 5h, i**). We did not observe changes in the number of ATAC fragments per cell between the different perturbed genes (**Supplementary Fig. 6a, b**) and gene enrichment was not correlated with the number of ATAC fragments, peaks, or differential peaks obtained from sgRNAs targeting the same gene (**Supplementary Fig. 6c-e**).

We examined how loss of these chromatin modifiers impacts accessibility within known chromatin marks (primarily histone post-translation modifications) using ENCODE ChIP-seq data from K562 cells (**Fig. 2a, Supplementary Tables 4, 5**). We found similar accessibility changes between different sgRNAs targeting the same genes, further highlighting the consistency between distinct genetic perturbations targeting the same gene (**Fig. 2b**). Targeting the Polycomb repressive complex 2 (PRC2) subunit EZH2 resulted in an increase in chromatin accessibility at regions with H3K27me3, a marker of heterochromatin (**Fig. 2a**). EZH2 catalyzes nucleosome compaction via H3K27 trimethylation^7^ and thus loss of EZH2 increases accessibility in these regions. Differential accessibility at known chromatin marks can be highly specific: A downsampling analysis reveals that for some target genes, like *EZH2* and *ARID1A*, a small number of cells correlates well (*r_p_* ≥ 0.75) to an aggregated (pseudo-bulk) cell population (5 single cells for *EZH2*, 25 single cells for *ARID1A*) (**Fig. 2c, Supplementary Fig. 7a, b**). For cells receiving a non-targeting sgRNA, we find that 75 cells correlate well with the respective pseudo-bulk populations (**Supplementary Fig. 7c**). Over all perturbations, we find that the median cell number to represent the pseudo-bulk is also 75 cells. For some CRISPR perturbations more cells are needed to accurately represent the pseudo-bulk (e.g. 225 cells for *TET2*) (**Supplementary Fig. 7d**), indicating that disruption of these genes creates more variable chromatin accessibility than non-targeting controls.

**Fig. 2.**
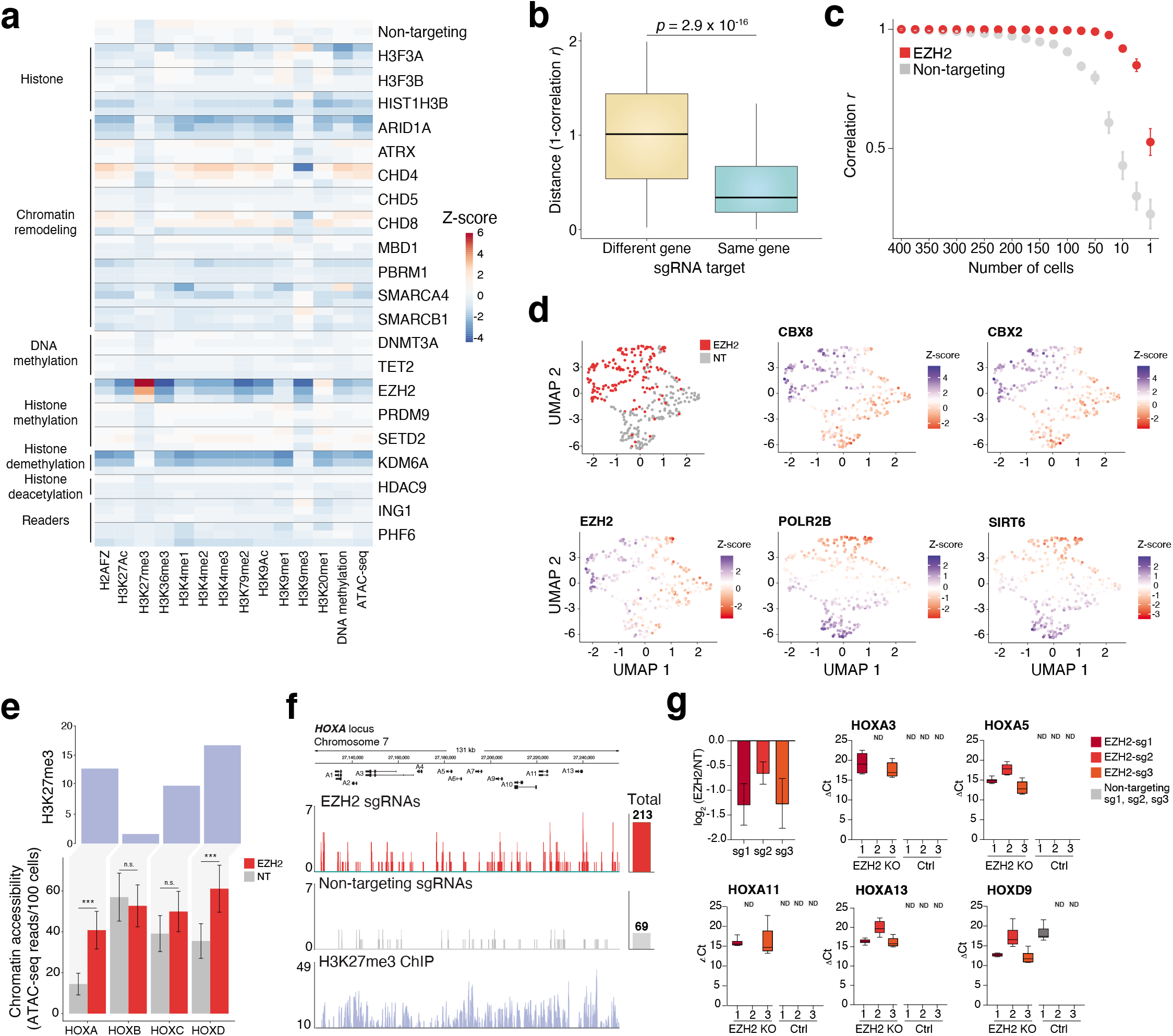
CRISPR-sciATAC reveals changes in accessibility at *HOX* genes following loss of EZH2. (**a**) Heatmap of chromatin accessibility *Z*-scores at histone and DNA modifications for different CRISPR perturbations (*n* = 3 sgRNA per gene). We converted the fraction of accessible regions for each modification into *Z*-scores (using all cells in the screen). For visualization, we show the average *Z*-score for all cells receiving a particular sgRNA. (**b**) Distances in the histone and DNA modifications accessibility profiles shown in panel *a* between sgRNAs targeting different genes and sgRNAs targeting the same gene. The distance metric is 1-(Pearson correlation of the *Z*-scores). (**c**) Pearson correlation between averaged accessibility *Z*-scores at histone and DNA modifications of the indicated number of single cells and the average profile of 400 single cells, for cells with either *EZH2*-targeting or non-targeting (NT) sgRNAs. (**d**) UMAP representation of chromatin accessibility *Z*-scores at histone and DNA modifications from single cells receiving either *EZH2* or NT sgRNAs. Also shown is the same UMAP representation with single cells colored by TFBS accessibility enrichment scores for *CBX2, CBX8, EZH2, POLR2B*, and *SIRT6*. (**e**) H3K27me3 ChIP-seq coverage at the *HOXA-D* loci (*top*). Changes in accessibility at the *HOXA-D* loci in cells transduced with *EZH2*-targeting or NT sgRNAs (*bottom*). *** denotes *p* < 0.001. (**f**) CRISPR-sciATAC fragments mapping to the *HOXA* locus in cells transduced with *EZH2*-targeting or NT sgRNAs (*n* = 510 cells per condition). The sum of all ATAC fragments over the entire *HOXA* locus in cells transduced with *EZH2*-targeting and NT sgRNAs is shown on the right. K562 H3K27me3 ChIP-seq coverage is shown at the bottom. (**g**) Gene expression (qPCR) of *EZH2, HOXA3, HOXA5, HOXA11A, HOXA13* and *HOXD9* for cells transduced with either *EZH2*-targeting or NT sgRNAs

Hierarchical clustering of single cells transduced with *EZH2*-targeting sgRNAs or non-targeting sgRNAs reveals a clear separation (**Supplementary Fig. 8a, b**), that can also be observed in a uniform manifold projection (UMAP) (**Fig. 2d**). We verified this separation is not due to differences in library complexity in cells with *EZH2*-targeting sgRNAs (**Supplementary Fig. 8c**), and found that increased accessibility in Polycomb repressive complex 1 (PRC1) components CBX2 and CBX8 binding sites has the highest predictive power in differentiating EZH2-targeted cells. Similarly, differential accessibility of POLR2B and SIRT6 binding sites can also be used to differentiate between cells with *EZH2*-targeting and non-targeting sgRNAs, however in the opposite direction, where loss of EZH2 leads to decreased accessibility in their binding sites. As expected, we find an increase in accessibility at EZH2 binding sites, which is expected given EZH2’s role in repression through heterochromatin formation^8^.

Using Gene Ontology (GO) analysis of differentially accessible regions in *EZH2*-targeted cells, we found an enrichment in genes involved in embryonic development and cell differentiation (**Supplementary Fig. 9, Supplementary Table 6**). Indeed, EZH2 is known to play important roles in embryonic development and cell- and tissue-specific differentiation^7^ and we found large changes in chromatin accessibility at several of the homeobox (HOX) genes (**Fig. 2e**). In K562 cells, the *HOXA* and *HOXD* gene clusters contain the highest amount of H3K27me3 repressive heterochromatin. In the *HOXA* gene cluster, we found that there was a nearly 3-fold increase in accessibility (**Fig. 2f**). A similar increase in accessibility was also seen at the *HOXD* gene cluster (**Supplementary Fig. 9d**). To understand the functional consequences of these changes, we measured the expression of *EZH2* and several HOX genes (*HOXA3, HOXA5, HOXA11, HOXA13*, and *HOXD9*) (**Fig. 2g**). After EZH2 loss, we found that these previously-silenced genes become highly expressed. Among the 3 sgRNAs targeting *EZH2*, we noticed that the least effective sgRNA resulted in the smallest increase in HOX gene expression, further reinforcing the role of EZH2 in maintaining HOX gene repression. Taken together, these results suggest that loss-of-function mutations in *EZH2* lead to aberrant expression of genes from *HOXA* and *HOXD* clusters.

In comparison to bulk ATAC-seq, an advantage of single-cell ATAC-seq is the ability to determine regulatory relationships between transcription factors (TFs) and their heterogeneity across cells. We wondered if loss of EZH2 might also impact co-regulation of particular TFs. To analyze this, we computed the correlation in accessibility between pairs of TFs before and after loss of EZH2. In cells receiving a non-targeting sgRNA, we found that ~90% of the 6,555 TF-TF pairs tested have some correlation in their accessibility (*r_p_* > 0.3) across single cells. However, upon EZH2 loss, a subset of these TF-TF correlations are disrupted (44 TF pairs with Δ*r_p_* < −0.3) (**Supplementary Fig. 10a**). Of these 44 TF pairs, we found that 40 of them include CREBBP, a ubiquitous coactivator of many different TFs, such as MYC and CEBPB, that couples chromatin remodeling to TF recognition^9^ (**Supplementary Fig. 10b,c**). This suggests that EZH2 may recruit CREBBP in a coordinated fashion with several other TFs. We found that these 40 TFs whose coregulation with CREBBP is disrupted are enriched in pathways central to transcriptional regulation in cancer and interact directly with other oncogenic master TFs like EP300 (**Supplementary Fig. 10d, e**).

Beyond EZH2, we found that changes in accessibility in single cells at transcription factor binding site (TFBS) are consistent between sgRNAs targeting the same gene (**Supplementary Fig. 11a, b**). We also noticed that large changes in TF accessibility correlate with decreased cell proliferation (**Supplementary Fig. 11c**), suggesting that perturbing chromatin modifiers which broadly disrupt TFBS can impact cell viability. We found similar changes in TFBS accessibility using either predicted TF binding sites (JASPAR TF motifs^10^ and chromVAR^11^) or ENCODE ChIP-seq data from K562 (**Supplementary Fig. 11d,e**). To determine if chromatin accessibility is modified at single nucleotide polymorphisms (SNPs) that regulate gene expression, we measured overlap with *cis*-regulatory expression quantitative trait loci (*cis*-eQTLs). For two of our targets — *KDM6A* and *ARID1A* — we found a reduction in accessibility at tissue-matched (blood) *cis*-eQTLs in cells after perturbation of these genes (**Supplementary Fig. 12a**). *KDM6A*-targeted cells had the largest reduction of *cis*-eQTL accessibility with eQTL genes (eGenes) involved in DNA condensation and chemokine receptor activity (**Supplementary Fig. 12b-e**).

To further demonstrate the scalability of CRISPR-sciATAC, we designed a CRISPR library to target all human chromatin remodeling complexes in the EpiFactors database^12^ (**Fig. 3a**). In total, we targeted 17 chromatin remodeling complexes that each include between 2 and 14 subunits. As before, we targeted the coding exons of each subunit with 3 sgRNAs and also included sgRNAs designed not to target anywhere in the human genome. Over the 17 chromatin remodeling complexes, we captured paired CRISPR perturbation and single-cell ATAC-seq data from 16,676 cells. As in the previous screen, the number of cells recovered for each CRISPR perturbation correlated with gene essentiality scores^6^ (**Supplementary Fig. 13a**). We recovered particularly low numbers of cells for the two subunits of the FACT complex, which are known to be highly essential^13^ (**Supplementary Fig. 13a, b**).

**Fig. 3.**
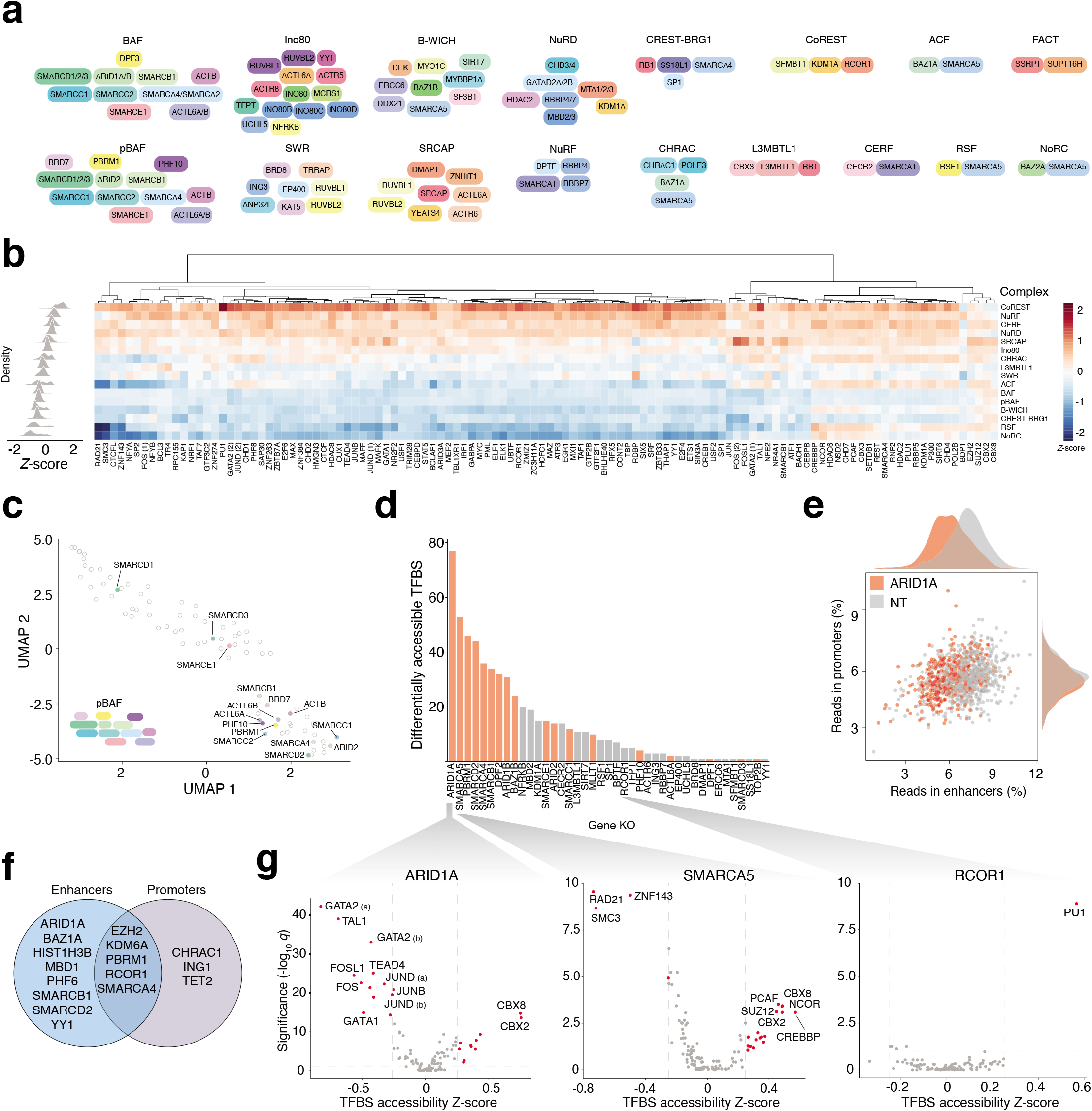
A CRISPR-sciATAC screen targeting 16 chromatin remodeling complexes uncovers widespread disruptions in accessibility upon SWI-SNF disruption. (**a**) Chromatin remodeling complex subunits and cofactors targeted in the CRISPR library. (**b**) Heatmap of chromatin accessibility *Z*-scores at transcription factor binding sites (TFBSs) for the different chromatin remodeling complexes targeted in the screen. We converted the fraction of accessible regions for each TFBS into *Z*-scores (using all cells in the screen). For visualization, we first average over all cells for a particular target gene and then average over all genes in the complex. The histograms (*left*) show the distribution of *Z*-scores for each complex. The FACT complex is not shown due to a low number of single cells (*n* = 75 cells). (**c**) UMAP representation of the genes perturbed in the screen based on the TFBS differential accessibility *Z*-score profiles. Subunits of the SWI-SNF pBAF complex are labeled with filled circles and gene names. (**d**) The number of transcription factor binding sites with significant differential accessibility for cells that receive a specific genetargeting CRISPR perturbation, as compared to cells that receive a non-targeting (NT) control sgRNA (FDR *q* ≤ 0.1). SWI/SNF components and co-factors are highlighted in red. (**e**) The percent of ATAC fragments in enhancers and promoters in cells transduced with *ARID1A*-targeting and NT sgRNAs. Each point is a single cell. K562 enhancer and promoter genome segmentation is from ENCODE (see *Methods*). (**f**) CRISPR-targeted chromatin complex genes with significant differential accessibility at enhancers and/or promoters. (**g**) Volcano plots showing significant changes in accessibility at TFBSs in cells transduced with *ARID1A (left), SMARCA5* (*middle*) and *RCOR1* (*right*) -targeting sgRNAs. Standardized *Z*-scores are averaged over single cells. Points in red represent TFBSs with a significant change in accessibility (FDR *q* ≤ 0.1 and |*Z*-score| > 0.25).

Given the larger scale of this CRISPR-sciATAC screen, we initially analyzed changes in accessibility at the level of different chromatin remodeling complexes instead of individual proteins/subunits (**Fig. 3b**). Examination of differential accessibility in TFBSs revealed two major groups: Complexes where loss of subunits generally results in increased accessibility, such as the CoRepressor for Element-1-Silencing Transcription factor (CoREST) and the Nucleosome Remodeling Factor (NuRF) complexes, and another group where loss of subunits leads to decreased accessibility, such as the Calcium RESponsive Transactivator-BRG1 (CREST-BRG1) and SWI/SNF-B (pBAF) complexes. However, loss of individual subunits within these complexes display tremendous heterogeneity: Nearly all complexes have subunits where loss triggers increased accessibility and other subunits with the opposite effect (**Supplementary Fig. 14, Supplementary Table 7**). A two-dimensional UMAP projection of the TFBS accessibility profiles reveals a cluster enriched in SWI/SNF components and, in particular, pBAF components (hypergeometric *p* = 4 x 10^−4^) (**Fig. 3c**). Loss of SWI/SNF subunits tends to alter accessibility at many TFBS, with the greatest number of disrupted TFBSs from ARID1A loss (**Fig. 3d**). Previously, ARID1A loss has been shown to impair enhancer-mediated gene regulation^14^, and indeed we find that loss of ARID1A dramatically reduced accessibility at enhancers, but not at promoters (**Fig. 3e**).

Combining data from both CRISPR-sciATAC experiments, we found that the chromatin modifiers targeted in our two screens resulted in a greater number of accessibility changes at enhancers than at promoters (**Fig. 3f**), supporting a gene regulatory model with more dynamic chromatin accessibility at distal regulatory elements compared to promoters^15^. Loss of SWI/SNF-ATPase subunit ARID1A or ISWI-ATPase subunit SMARCA5 results in many changes in TFBS accessibility (**Fig. 3g**). For ARID1A, some of these changes include a reduction in accessibility at JUN and FOS binding sites, which are subunits of the AP-1 transcription factor that cooperate with the SWI/SNF complex to regulate enhancer activity^16^. Loss of SMARCA5, which helps load cohesion onto chromosomes^17^, triggered a reduction in accessibility in binding sites of cohesin subunits RAD21 and SMC3 along with cohesin cofactor ZNF143^18^. In contrast to these gene perturbations that affect a wide range of TFBSs, other perturbations result in accessibility changes at only one or a few TFBSs. For example, we observe an increase in accessibility at only PU.1 binding sites upon loss of RCOR1 (**Fig. 3g**). RCOR1 has previously been shown to promote erythroid differentiation via repression of myeloid genes such as PU.1 and thus may have a focused role in lineage specification^19^.

In addition to changes in accessibility, chromatin remodeling complexes can regulate gene expression by changing specific nucleosome positions around regulatory sequences^20^. We developed a computational framework to measure changes in nucleosome position in CRISPR-sciATAC for TFs with symmetric positioning of nucleosomes around their binding sites^21^ (**Fig. 4a, b**). Using this pipeline, we found that loss of chromatin remodelers generally results in expansion of nucleosomes around TFBSs (**Fig. 4c**), with the exception of BAF/pBAF (SWI/SNF) subunits ARID1A and PBRM1 where knock-out leads to compaction of nucleosomes around the TFBSs studied (**Fig. 4b**). At specific TFBS, loss of different chromatin remodelers can have opposing effects: For example, ARID1A loss results in a 20 nt nucleosome compaction at AP-1 binding sites (*p* = 0.03), which has also been demonstrated in a recent study suggesting that the BAF complex controls occupancy of AP-1^22^. In contrast, loss of EP400, which is part of the Sick With Rat8ts (SWR) complex, causes a large, 56 nt expansion of nucleosomes around AP-1 binding sites (*p* = 10^−4^) (**Fig. 4d**).

**Fig. 4.**
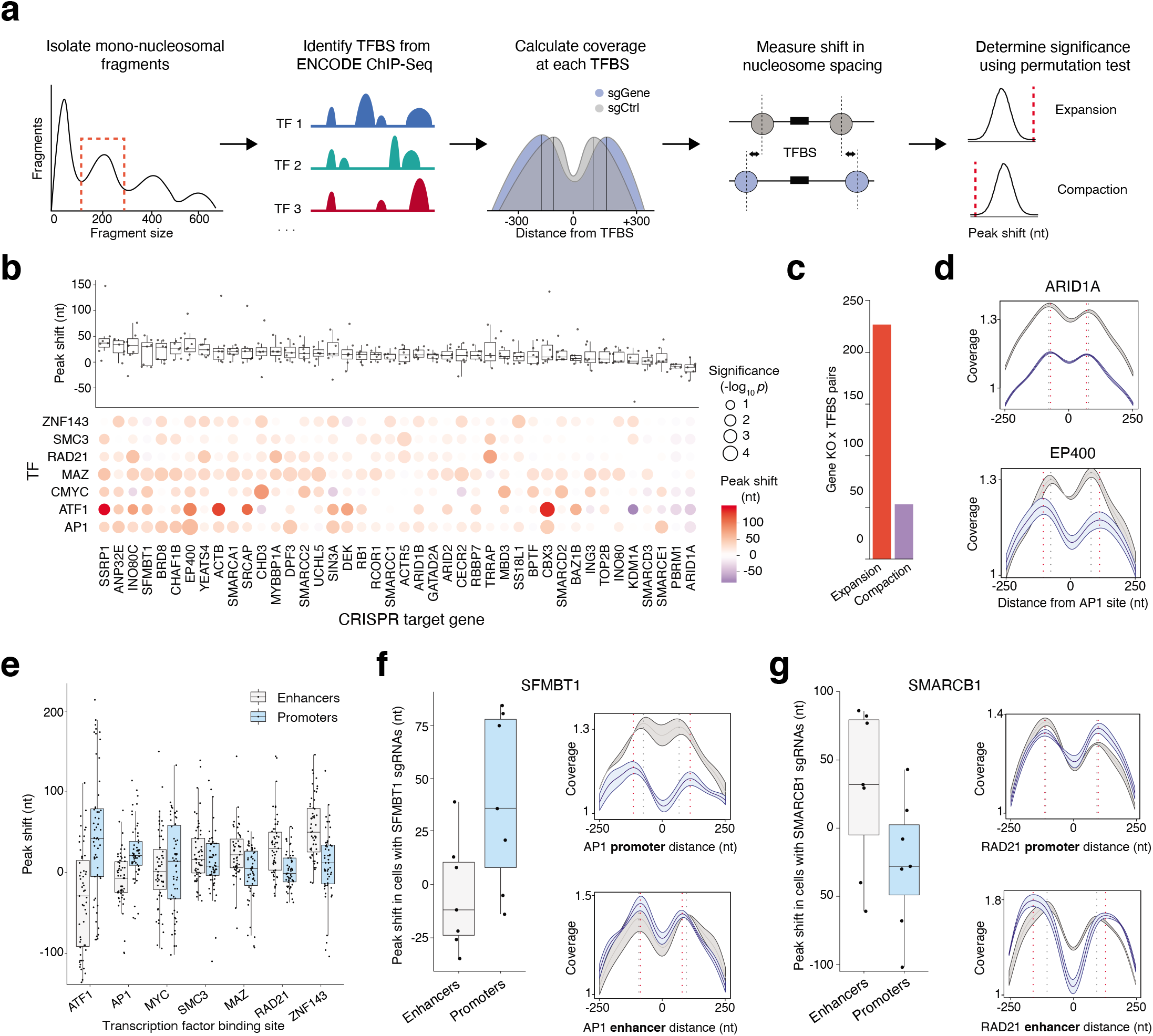
Nucleosome dynamics around transcription factor binding sites (TFBSs) following CRISPR targeting of chromatin remodelers. (**a**) Schematic depicting the computational approach to identify changes in nucleosome positions around TFBSs. (**b**) Absolute peak shift across 7 TFBS following CRISPR targeting of chromatin remodelers (*top*). Bubble-plot of the peak shifts for individual TFBS (*bottom*). The color of the bubble corresponds to the peak shift (nt) and the size of the bubble represents the empirical *p*-value calculated by a label permutation test. (**c**) The number of nucleosome expansion and compaction events around TFBSs following CRISPR targeting of chromatin remodelers. (**d**) Coverage profiles of mono-nucleosomal fragments around AP1 binding sites in cells transduced with *ARID1A*-targeting (blue) and nontargeting (NT) (grey) sgRNAs (*top*) and in cells transduced with *EP400-targeting* (blue) and NT (grey) sgRNAs (*bottom*). Dashed lines represent the most highly covered base in each peak. (**e**) Peak shifts in TFBSs located in enhancers and promoters. Each point is a CRISPR-perturbed gene (average of all sgRNAs for that gene). (**f**) Peak shifts in TFBSs located in enhancers and promoters in *SFMBT1*-targeted cells (*left*). Coverage profiles of mono-nucleosomal fragments in cells transduced with *SFMBT1*-targeting (blue) and NT (grey) sgRNAs around AP1 binding sites in promoters (*top*) and in enhancers (*bottom*). (**g**) Peak shifts in TFBSs located in enhancers and promoter in *SMARCB1* targeted cells (*left*). Coverage profiles of mono-nucleosomal fragments in cells transduced with *SMARCB1*-targeting (blue) and NT (grey) sgRNAs around RAD21 binding sites in promoters (*top*) and in enhancers (*bottom*). For panels *d*, *f*, and *g*, the shaded regions represent s.e.m. (*n* = 3 sgRNAs).

We further asked if there are specific differences in nucleosome dynamics surrounding TFBSs residing in enhancers versus promoters. We found that changes in nucleosome peak positions occur typically in either enhancers or promoters, depending on the specific TFBS. For example, across all CRISPR perturbations, the expansion of nucleosome spacing around AP-1 binding sites occurs mostly in sites that are located in promoters (**Fig. 4e**). In contrast, expansion of nucleosome spacing around ZNF143 binding sites occurs mostly in sites that are located in enhancers. An exception to this trend is ATF1: Knock-out of chromatin remodelers results in nucleosome expansion around ATF1 binding sites in promoters, but compaction in ATF1 binding sites in enhancers (**Supplementary Fig. 15a, b**). For specific chromatin modifiers, we often observed more expansion in either enhancers or promoters (**Supplementary Fig. 15c**). Knock-out of CoREST subunit SFMBT1 tends to cause nucleosome expansion around TFBSs in promoters but not in enhancers: for example, an 85 nt expansion around AP-1 binding sites in promoters and no change in nucleosomal positions around AP-1 binding sites in enhancers (**Fig. 4f**). In contrast, knock-out of BAF/pBAF subunit SMARCB1 tends to cause nucleosome expansion around TFBSs in enhancers but not in promoters (e.g. at RAD21 sites) (**Fig. 4g**).

In this work, we develop CRISPR-sciATAC, a platform for pooled forward genetic screens that jointly captures CRISPR perturbations and ATAC profiles in single cells. Pooled CRISPR screens have been used extensively to identify genes responsible for therapeutic resistance, cell proliferation, and Mendelian disorders^23^. In recent years, CRISPR screens have been combined with single-cell RNA-sequencing to measure the effects of genetic perturbations on gene expression across the transcriptome^3,24–26^. However, methods to capture changes in the epigenome following CRISPR perturbations have been limited. Rubin and collaborators^27^ published a related method (Perturb-ATAC) which uses a programmable microfluidic device to physically isolate single cells into small chambers. This method delivers high-depth single-cell ATAC-seq data (~10^4^ fragments per cell), but the throughput per experiment is limited to the 96 chambers of the microfluidic device. CRISPR-sciATAC offers an alternative approach that takes advantage of two-step combinatorial indexing to label DNA molecules with unique cell barcodes and requires no specialized equipment. When compared with Perturb-ATAC, CRISPR-sciATAC can generate thousands of single cells at ~20x less reagent cost and requires ~14x less time (**Supplementary Tables 8 and 9**). In this work, we analyzed 28,510 cells, which is ~7-fold more cells than in the Perturb-ATAC dataset (**Supplementary Fig. 16a**). Using a library of 318 sgRNAs targeting 105 genes, we investigated differential accessibility at histone and DNA modifications and at TFBSs following loss of chromatin modifiers. By perturbing chromatin remodeling complexes in a high-throughput and uniform setting, we reduce batch effects and generate data for a large number of different chromatin complexes. Since it is based on combinatorial indexing ATAC-seq, CRISPR-sciATAC shows comparable yield to published sciATAC datasets, which do not have the additional modality of sgRNA capture (**Supplementary Fig. 16b**). As we demonstrate with sgRNAs targeting *EZH2*, one important caveat of CRISPR nuclease-driven perturbation is that knock-out can be incomplete due to in-frame repair and efficiency can vary depending on the guide RNA. However, CRISPR-sciATAC does not depend critically on any specific guide RNA but rather looks for consistent effects between guide RNAs. To more completely address these issues, future computational methods to discern perturbed cells from unperturbed cells, as has been recently developed for single-cell RNA-sequencing^28^, will also be useful for single-cell ATAC-seq. Overall, CRISPR-sciATAC can be applied to study diverse phenotypes and diseases and to understand the interaction between genetic changes and genome-wide chromatin accessibility.

## Supporting information

Supplementary Figures and Sequences

## Acknowledgements

We thank the entire Sanjana laboratory for support and advice. We also thank J. Morris for help with eQTL resources, M. Zaran and R. Satija for computational resources and the NYGC Sequencing Platform and NYU Biology Genomics Core for sequencing resources. BL21(DE3) cells transformed with pET-PfuX7 were kindly provided by J. Gregory. N.L.B is supported by a postdoctoral fellowship from the Human Frontier Science Program Organization (LT000672/2019-L), an EMBO long-term fellowship (ALTF 826-2018), and the Weizmann Institute of Science National Postdoctoral Award Program for Advancing Women in Science. N.E.S. is supported by NYU and NYGC startup funds, NIH/NHGRI (R00HG008171, DP2HG010099), NIH/NCI (R01CA218668), DARPA (D18AP00053), the Sidney Kimmel Foundation, the Melanoma Research Alliance, and the Brain and Behavior Foundation.

## Authors contributions

N.E.S. conceived and supervised the project. N.E.S., A.M. and N.L.B. designed the experiments. A.M., N.L.B., J.D., A.M.M., C.Y.K., and A.S. performed the experiments. N.L.B., A.M., J.D., N.E.S., H.H.W., and N.G.M. analyzed the data. P.S. isolated TnY. S.J. purified PhuX7. A.M., J.D., C.Y.K., A.S., P.S. and S.J. purified TnY. N.L.B., A.M. and N.E.S. wrote the manuscript with input from all of the authors.

## Competing interests

The New York Genome Center and New York University have applied for patents relating to the work in this article. N.E.S. is an adviser to Vertex.

## Methods

### Cell culture and monoclonal K562-Cas9 cell line

NIH-3T3 and K562 cells were acquired from ATCC (CRL-1658 and CCL-243). HEK293FT cells were acquired from Thermo Fisher (R70007). NIH-3T3 (mouse) and HEK293FT (human) cells were maintained at 37°C with 5% CO_2_ in D10 media: DMEM with high glucose and stabilized L-glutamine (Caisson DML23) supplemented with 10% fetal bovine serum (Thermo Fisher 16000044). K562 cells were maintained at 37°C with 5% CO_2_ in R10 media: RPMI with stabilized L-glutamine (Thermo Fisher 11875119) supplemented with 10% fetal bovine serum. To generate monoclonal K562 cells expressing Cas9, K562 cells were transduced with lentiCas9-Blast (Addgene 52962) at a multiplicity of infection (MOI) of 0.1 and selected and maintained in R10 with 5 μg/ml blasticidin. Monoclonal K562-Cas9 cells were isolated and expanded through limiting dilution. Expression of Cas9 was confirmed by Western blot using an anti-2A peptide antibody (Millipore Sigma MABS2005).

### Lentiviral CRISPR libraries

To generate NIH-3T3 and HEK293FT cells expressing single-guide RNAs (sgRNAs) for the human/mouse experiment, 10 human non-targeting sgRNAs and 10 mouse non-targeting sgRNAs (**Supplementary Table 2**) were individually synthesized and cloned into the lentiviral transfer vector CROPseq-Guide-Puro^3^ (Addgene 86708), which leads to the synthesis of an RNA Pol3 transcript of the Cas9 sgRNA and an RNA Pol2 polyadenylated transcript containing the puromycin resistance gene, a U6 promoter, and the sgRNA. The RNA Pol2 transcript allows for the selection of transduced cells (via puromycin) and detection of the sgRNA targeting sequence via reverse-transcription and PCR. During reverse-transcription, our priming strategy (using a primer that anneals to the sgRNA scaffold) can capture either the sgRNA encoded at the 3’ end of the Pol2 transcript (**Supplementary Fig. 1c**) or the sgRNA in the Pol3 transcript. Equal amounts of each sgRNA plasmid were mixed and then, with packaging plasmids pMD2.G (Addgene 12259) and psPAX2 (Addgene 12260), transfected into HEK293FT cells^29^. NIH-3T3 and HEK293FT cells were transduced at MOI ~ 0.1 and selected and maintained in D10 with 1 μg/ml puromycin. The sgRNA library coverage is 1,500X on average for the species-mixing experiment, the chromatin modifier screen and the chromatin remodeling complex subunit screen.

For the chromatin modifier pooled CRISPR screen, we identified 21 frequently mutated chromatin modifiers across all cancers in the Catalogue of Somatic Mutations in Cancer (COSMIC) database^4^ (**Supplementary Fig. 4a,b**) and designed three targeting sgRNAs per gene. An important issue to consider when designing sgRNAs is that not all CRISPR nuclease-driven modifications will result in loss-of-function, as some genome modifications (non-homologous end-joining) will result in in-frame repair that may preserve gene function. However, it has been previously demonstrated that even in-frame mutations can be disruptive when targeting functional domains of proteins^30^. To capitalize on this discovery, we chose a CRISPR library design algorithm that uses protein functional domains (from the Pfam database) to target our guide RNAs^31^. The final library was composed of 63 targeting and 3 non-targeting sgRNAs that were individually synthesized (IDT) and annealed (**Supplementary Table 2**). For the chromatin remodeling complex subunit pooled CRISPR screen, we designed a CRISPR library to target all chromatin remodeling complexes in the human genome, as defined by the EpiFactors database^12^ (**Fig. 3a**). The library was composed of 252 targeting and 3 non-targeting sgRNAs that were individually synthesized (IDT) and annealed (**Supplementary Table 2**). Annealed oligos were pooled in equimolar ratio and cloned as a pool into the CROPseq-Guide-Puro lentiviral transfer vector. K562-Cas9 cells were transduced at a MOI of ~0.1 and selected and maintained in 1 μg/ml puromycin and 5 μg/ml blasticidin. The CRISPR-sciATAC protocol was performed on these cells at one week postselection.

### Transposase identification and isolation

We were motivated to use a different transposase than Tn5 due to the difficulty of obtaining sufficient yields of Tn5^32^. In order to identify new transposases, sequences were aligned using ClustalW^33^. We found a range of transposon sequences that were related to the Tn5 sequence and selected a transposon from *Vibrio parahemolyticus* (ViPar) for further analysis. The inside and outside ends (IE and OE) of the ViPar transposon utilize the same sequence as the IE and OE of the Tn5 transposon, giving us confidence the ViPar transposon would be compatible with existing Tn5-based workflows (**Supplementary Fig. 2a and b**). The identified ViPar transposase was synthesized (Twist Bioscience) and cloned into the vector pTXB1 (NEB, N6707S). Two mutations were introduced: (1) P50K, equivalent to the mutation E54K in Tn5, which is predicted to make the transposon hyperactive^34^ and (2) M53Q, which changes the residue that interacts with nucleotide 9 (a thymine) on the non-transferred strand of the mosaic end (ME) similar to Tn5 Q57, predicted to increase binding to the Tn5 ME. The ViPar transposase with P50K and M53Q mutations, henceforth referred to as TnY, showed Tn5 ME loading and tagmentation activity (**Supplementary Fig. 2c-f**). Finally, we characterized the insertion site preference of TnY by performing tagmentation on NA12878 DNA and sequencing on a MiSeq Instrument (Illumina); we found that TnY has insertion site preferences distinct from, but of a similar magnitude to those of Tn5 (**Supplementary Fig. 2g, h**). The chromatin accessibility profiles resulting from TnY and Tn5 are highly correlated (**Supplementary Fig. 2i, j**).

### TnY transposase production

The pTXB1-TnY vector was transformed into BL21(DE3) competent *E. coli* cells (NEB C2527) and TnY was produced via intein purification with an affinity chitin-binding tag^32^. One liter of LB culture was grown at 37°C to OD600 = 0.6. TnY expression was then induced with IPTG 0.5 mM at 18°C overnight. After induction, cells were pelleted and then frozen at −80°C overnight. Cells were then lysed by sonication in 100 ml HEGX (20 mM HEPES-KOH at pH 7.5, 0.8 M NaCl, 1 mM EDTA, 10% glycerol, 0.2% Triton X-100) with a protease inhibitor cocktail (Roche 04693132001). The lysate was pelleted at 30,000 x g for 20 min at 4°C. Supernatant was transferred to a new tube, 3 μl of neutralized PEI 8.5% (Sigma Aldrich P3143) was added dropwise to each 100 μl of bacteria extract, gently mixed and centrifuged at 30,000 x g for 30 minutes at 4°C to precipitate DNA. The supernatant was loaded on four 1-ml chitin columns (NEB S6651S). Columns were washed with 10 ml HEGX; 1.5 ml HEGX containing 100 mM DTT was added to the column and incubated for 48 h at 4°C to allow cleavage of TnY from the intein tag. TnY was eluted directly into two 30 kDa MWCO spin columns (Millipore UFC903008) by adding 2 ml of HEGX. Protein was dialyzed in five dialysis steps using 15 ml 2x Dialysis Buffer (100 HEPES-KOH at pH 7.2, 0.2 M NaCl, 0.2 mM EDTA, 2 mM DTT, 20% glycerol) and concentrated to 1 ml by centrifuging at 5,000 x g. The protein concentrate was transferred to a new tube and mixed with an equal volume of glycerol 100%. Then, we added Triton X-100 (0.04% final concentration). TnY aliquots were stored at −80°C.

### Transposome assembly

To produce mosaic-end double-stranded (MEDS) oligos, we annealed the single T5 tagmentation oligo with the pMENT common oligo (100 μM each) (**Supplementary Table 2**) as follows in TE buffer: 95°C for 5 minutes, then cooled at a rate of 0.2°C /s down to 4°C (“MEDS A”). The same process was used to anneal each barcoded T7 tagment sciATAC oligo with the pMENT common oligo (“MEDS B”) (**Supplementary Table 2**). MEDS A and MEDS B were mixed together, diluted 1:6 in TE buffer and 2 μl were transferred into a new tube and mixed with 3 μl of TnY enzyme. After 30 minutes at room temperature to allow for transposome assembly, we added 45 μl Dilution Buffer, mixed by pipetting up and down and stored at −20°C until ready for tagmentation. Dilution Buffer consists of 2x Dialysis Buffer (see *TnY transposase production* above) diluted 1:1 by volume with 100% glycerol. We observed optimal tagmentation when transposome assembly was carried out on the same day as the CRISPR-sciATAC tagmentation.

### PfuX7 polymerase production

For CRISPR-sciATAC, we used a purified PfuX7 DNA polymerase^35^. First, we transformed BL21(DE3) competent *E. coli* cells (NEB C2527) with pET-PfuX7 and grew them in 1 L of LB culture at 37°C to OD_600_ = 0.6. PfuX7 expression was then induced with IPTG (0.5 mM final concentration) at 30°C overnight. After induction, cells were pelleted and resuspended in 20 ml Lysis Buffer (50 mM Tris-HCl pH8, 150 mM NaCl, 1 mM EDTA, 1 mM PMSF, 10 μg/ml EDTA-free protease inhibitor (Sigma 11873580001)) and sonicated in an ice slurry. Sonication was at 20% amplitude for ten cycles of 1 minute duration with a 30 second pause between cycles (Branson Ultrasonics, Model 450 Digital Sonifier). The lysate was pelleted at 30,000 x g for 15 min at 4°C. Supernatant was transferred to a new tube and incubated with DNA Digestion Buffer (20 μl DNaseI (NEB M0303), 0.5 mM CaCl_2_, 2.5 mM MgCl_2_) for 30 minutes at 37°C. DNaseI was then inactivated by incubating for 30 minutes at 85°C. After inactivation, the lysate was placed on ice for 20 minutes. Lysate was then centrifuged at 50,000 x g for 20 minutes at 4°C. Supernatant was loaded on two 1-ml Ni-NTA (Qiagen 30210) columns, washed twice with Wash Buffer (50 mM Tris-HCl pH 8, 150 mM NaCl). PfuX7 enzyme was eluted in 5 ml Elution Buffer (50 mM Tris-HCl pH 8, 150 mM NaCl, 0.25 M imidazole) and desalted in Storage Buffer (100 mM Tris-HCl pH 8, 0.2 mM EDTA, 2 mM DTT) by performing buffer exchange three times using one Amicon 30 kDa MWCO spin column (Millipore UFC903008). The purified protein was then transferred to a new tube, combined with equal volume of 100% glycerol and adjusted with Tween-20 (0.1% final concentration) and IGEPAL CA630 (0.1% final concentration). Aliquots were stored at −20°C.

### Bulk ATAC-seq

For bulk ATAC-seq^36^, we resuspended 500,000 cells in 1 ml PBS and gently lysed them by adding 10 ml Resuspension Buffer (10 mM Tris-HCl at pH 7.5, 10 mM NaCl, 3 mM MgCl_2_) with 0.1% Tween-20. Cells were then centrifuged at 500 *xg* for 10 min at 4°C to pellet the nuclei. Pelleted nuclei were resuspended in 600 μl 1x Tagmentation Buffer (10 mM TAPS-NaOH at pH 8.5, 5 mM MgCl_2_, 10% DMF), 30 μl (~25,000 nuclei) were then transferred into 1.5 ml tubes and 20 μl TnY transposomes were added. Tagmentation was performed at 37°C for 30 min. Samples were then purified using the DNA Clean & Concentrator kit (Zymo Research D4014) and eluted in 10 μl TE. Eluted DNA was thermocycled with PfuX7 in Phusion GC Buffer (Thermo Fisher F519L) as follows: 72°C 5 min, 98°C 30 s, (98°C 10 s, 63°C 30 s, 72°C 3 min) x 10 cycles, 4°C hold. Samples were purified using the DNA Clean & Concentrator kit, eluted in 6 μl TE and size-selected using a 0.9X volume of Ampure XP Beads (Beckman Coulter A63882) to remove excess oligos.

### CRISPR-sciATAC species mixing experiment in HEK293 and NIH3T3 cells

HEK293FT (human) and NIH-3T3 (mouse) transduced with non-targeting sgRNAs libraries were grown separately. On the day of the experiment, cells were counted, and 500,000 cells were resuspended in 1 ml PBS per cell line (1:1 ratio of human and mouse cells). Cells were then pelleted, resuspended in Fixation Buffer and fixed for 7 min at room temperature. Fixation Buffer consists of 2.8 ml H_2_O, 790 μl 100% ethanol, 310 μl 40% glyoxal (Sigma 128465), 30 μl glacial acetic acid (Sigma A6283); after preparing Fixation Buffer, adjust the pH to 5.0 by adding NaOH and keep ice-cold until immediately before use. In line with a previous study^37^, we found that glyoxal fixation resulted in better preservation of intact nuclei than the more commonly used paraformaldehyde fixative.

After fixation, cells were then washed three times with 1 ml PBS and gently lysed by adding and resuspending in 10 ml Resuspension Buffer (see *Bulk ATAC-seq* above) with 0.1% Tween-20 and 0.1% Igepal CA630. Cells were then incubated on ice for 3 minutes and then pelleted at 500 x*g* for 10 min at 4°C to obtain nuclei. Nuclei were washed in 1 ml Tagmentation Buffer (see *Bulk ATAC-seq* above) with 5 μl RiboLock RNase Inhibitor (ThermoFisher EO0381) and centrifuged at 500 x*g* for 5 min at 4°C. Human and mouse nuclei were resuspended and mixed together in a final volume of 3.2 ml Tagmentation Buffer with 28 μl RiboLock RNase Inhibitor. Nuclei (30 μl, ~20,000) were distributed into each well of a 96-well plate containing 20 μl of TnY assembled with MEDS A and 96 barcoded MEDS B (see **Supplementary Table 2** for MEDS sequences). Tagmentation was performed for 30 minutes at 37°C and then stopped by adding 2 μl EDTA 500 mM into each well. After incubating for 15 minutes at 37°C, EDTA was quenched prior to reverse transcription by adding 2 μl of 50 mM MgCl_2_ into each well.

For reverse transcription, 5 μl of the nuclei solution (~2,000 nuclei) were transferred into a new 96-well plate containing barcoded reverse transcription primers. Reverse transcription primers contain the same barcode as the MEDS B oligos (see **Supplementary Table 2** for RT oligos). Nuclei were transferred keeping plate orientation to match tagmentation and reverse transcription barcodes. The reverse transcription master mix (RTMM) consisted of 1 mL 5x RT buffer, 270 μl dNTPs, 1.6 mL water, 262 μl RevertAid reverse transcriptase, 27 μl RiboLock RNase Inhibitor (all components: Thermo Fisher, EP0442). We distributed 15 μl of RTMM into each well, mixed, and incubated for 30 min at 37°C.

Reverse transcription was stopped by adding 2 μl of Stop and Stain buffer (1 mL 500 mM EDTA, 2 μl of 5 mg/ml DAPI) and incubated for 5 minutes on ice. Nuclei were pooled together and pelleted at 500 x*g* for 5 min at 4°C. Supernatant was carefully removed taking care to not disturb the pellet. The nuclei were gently resuspended in 250 μl PBS and counted using a hemocytometer. PBS was added in order to obtain a final concentration of 10 nuclei/μl. 2 μl of the nuclei solution (~20 nuclei) were transferred into a new 96-well plate with DNA extraction and digestion buffer in each well. Specifically, each well contained 24.5 μl of DNA Rapid Extract Buffer (1 mM CaCl_2_, 3 mM MgCl_2_, 1% Triton X-100, 10 mM Tris-HCl at pH 7.5) and 2 μl of Digestion Buffer (1 μl H2O, 0.5 μl SDS 5.8%, 0.5 μl Proteinase K 20 mg/ml (Sigma P2308)). Nuclei were digested for 5 min at 65°C; digestion was stopped by adding 3 μl PMSF (Sigma 93482) and incubating for 30 min at room temperature.

For the first PCR, ATAC-seq primers and sgRNA-PCR1 primers were added at a final concentration of 0.5 μM and 0.1 μM, respectively. Amplification for ATAC-seq/sgRNA-PCR1 was performed with PfuX7 in Phusion GC Buffer as follows: 72°C 5 min, 98°C 30 s, (98°C 10 s, 63 °C 30 s, 72°C 3 min) x 14-18 cycles, 4°C hold. For the second PCR, 2 μl of PCR product were transferred into a new 96-well plate keeping plate orientation to match ATAC-seq and sgRNA barcodes. sgRNA-PCR2 primers were added to a final concentration of 0.5 μM. Amplification for sgRNA-PCR2 using PfuX7 in Phusion GC buffer was: 98°C 30 s, (98°C 10 s, 55°C 10 s, 72°C 20 s) x 20 cycles, 72°C 5 min, 4°C hold.

We then purified ATAC-seq and sgRNA amplicons. The ATAC-seq/sgRNA-PCR1 PCR plate was purified using four columns of the DNA Clean & Concentrator kit, eluted in 10 μl elution buffer and size-selected using 0.9X volume of Ampure XP Beads. The sgRNA-PCR2 PCR plate was purified using ten columns of the DNA Clean & Concentrator kit, eluted in 20 μl elution buffer. Eluted samples were run on E-gel 2% (Thermo Fisher G402002) and the expected band (~250 bp) gel extracted, purified using 1 column of Zymoclean Gel DNA Recovery Kit (Zymo Research D4008) and eluted in 20μl. Libraries were separately sequenced on the MiSeq Sequencer (Illumina) using the read lengths shown in **Supplementary Fig. 1d,e**^38,39^.

### CRISPR-sciATAC for chromatin modifiers in K562 cells

The CRISPR-sciATAC protocol for the chromatin modifier library in K562 cells was performed similarly to the human/mouse experiment described above. K562-Cas9 cells transduced with the pool of 63 chromatin modifiers sgRNAs and 3 non-targeting sgRNAs (library 1) or with the pool of 252 chromatin modifiers sgRNAs and 3 non-targeting sgRNAs (library 2) and were cultured for one week after selection. We prepared between either 12 (library 1) or 41 (library 2) 96-well plates and pooled amplicons. The ATAC-seq amplicons were sequenced on a HiSeq 2500 (Illumina) and the sgRNA amplicons were sequenced on a MiSeq.

### Gene essentiality screen and analyses

K562-Cas9 cells were transduced with the chromatin modifiers pooled CRISPR screen at MOI ~ 0.1 and selected and maintained in 1 μg/ml puromycin and 5 μg/ml blasticidin. Genomic DNA was extracted at three days (early time point), one week and two weeks post-selection. The sgRNA cassette was PCR amplified^2^. Libraries were sequenced on a MiSeq sequencer (Illumina). In addition to the CRISPR-sciATAC experiment, two independent transduction replicates were also analyzed. To identify essential genes, a *p*-value per sgRNA was calculated using the MAGeCK algorithm and *p*-values for the three sgRNAs targeting one gene were aggregated into a gene-level *p*-value using a Robust Rank Aggregation approach followed by a Bonferroni correction^5,40^.

### Read alignment

CRISPR-sciATAC sgRNA and ATAC datasets were demultiplexed based on cellular barcodes using the `snATAC_mat.py` script in an established sci-ATAC-seq pipeline (https://github.com/r3fang/snATAC)^41^. The processed sgRNA sequences were aligned to a custom guide reference using bowtie^42^ using the command `bowtie -v 1 -m 1`. For the human-mouse experiment, we show data for cells with at least 20 mapped reads. Cells with over 90% of sgRNA reads that mapped exclusively to human or mouse sgRNAs were considered species-specific cells. Cells where one sgRNA represented at least 90% of the total reads were kept for further analyses. The remaining cells were considered collisions and/or the result of multiple infections. For downstream analysis of the K562 data, we required each cell to have at least 100 aligned sgRNA reads with ≥ 99% of the reads assigned to one sgRNA sequence for the chromatin modifier screen and at least 10 aligned sgRNA reads with ≥90% of the reads assigned to one sgRNA sequence for the chromatin remodeling complex subunit screen. Gene knock-outs with at least 50 identified single cells were considered for further analysis (98/105 targeted genes).

The processed ATAC sequences were aligned to the reference genome using `bowtie2`^43^ using the command `bowtie2 -D 15 -R 2 -L 22 -i S,1,1.15 -p 5 -t -X2000 -e 75 --no-mixed --no-discordant`. The reference genome was a chimeric human hg19 and mouse mm10 genome for the human-mouse experiment and a human hg19 for the K562 datasets. Improperly paired and non-uniquely mapped alignments and reads mapping to mitochondrial DNA were removed. Reads overlapping ENCODE blacklist regions were removed (https://www.encodeproject.org/annotations/ENCSR636HFF/). Reads were then deduplicated using Picard (http://broadinstitute.github.io/picard). For the human-mouse experiment, we show data for cells with at least 20 unique ATAC-seq reads. For the K562 datasets, we require at least 500 unique ATAC-seq reads.

### Differential accessibility at genomic regions with specific chromatin and DNA modifications

To assess changes in accessibility, we downloaded from ENCODE ChIP-seq files covering post-translational histone modifications and DNA methylation (**Supplementary Table 4**). For each ChIP-seq track, we considered the fraction of fragments in each single cell that overlap ChIP-seq peaks. We standardized the averaged fractions over all single cells into *Z*-scores and then averaged the *Z*-scores obtained for each ChIP-seq file over cells that received the same sgRNA for the visualization in **Fig. 2a**. To find significant deviations in accessibility per gene-KO and per modification, we performed a two-tailed *t*-test on the *Z*-scores, of all cells for one gene knock-out and all the non-targeting cells, for each modification. The *p*-values were adjusted for multiple hypothesis testing using a Benjamini-Hochberg false-discovery rate correction (*q* ≤ 0.1).

For correlation of downsampled cell populations with the aggregated (pseudo-bulk) data, we randomly sampled cells (from a total of 400 single cells) without replacement. We performed this resampling procedure 200 times for each cell number. For each cell sample, we average the accessibility Z-scores and then compute the Pearson correlation with the pseudo-bulk. For this analysis, we only included target genes with at least 400 single cells.

### Differential accessibility in TF binding sites using ENCODE ChIP-seq

To identify enrichment or depletion in accessibility of TF binding sites following chromatin modifier knock-out, we downloaded 116 TF K562 ChIP-seq peak files from ENCODE (**Supplementary Table 4**) and considered the fraction of fragments in each single cell that overlap ChIP-seq peaks. We standardized the averaged fractions over all single cells into *Z*-scores and then averaged the *Z*-scores obtained for each ChIP-seq file over cells that received the same sgRNA for the visualization in **Supplementary Fig. 11a**. For dimensionality reduction, we used the function `umap` (from the R package `umap`) and, to predict cell perturbation, we fit TFBS *Z*-scores with a generalized linear model using the function `glm` (from the R package `stats`). To find significant deviations in accessibility per gene-KO and per TF, we performed a two-tailed *t*-test on the *Z*-scores, of all cells for one gene knock-out and all the non-targeting cells, for each TF. The *p*-values were adjusted for multiple hypothesis testing using a Benjamini-Hochberg false-discovery rate correction (*q* ≤ 0.1). For genes with multiple ENCODE ChIP-seq datasets, we denote with (1) ENCODE ChIP-seq profiles obtained using an antibody that directly recognizes the protein of interest; we denote with (2) ENCODE ChIP-seq profiles obtained using an antibody directed against an EGFP-tag.

### Differential accessibility in TF binding sites using JASPAR motifs

As an orthogonal method to ENCODE ChIP data, we also utilized predicted TF binding sites from the JASPAR database (386 motifs from JASPAR 2016, human CORE dataset)^10^. Transcription factor motif enrichment and depletion scores were calculated using chromVAR^11^. Briefly, Z-scores quantifying deviations in the frequency of each motif in each of the single cells were calculated based on the frequency of the motif in the collection of peaks that exist in each cell, out of all 358,028 peaks called on the aggregated single cell alignment files (pseudo-bulk). This frequency was compared to the frequency of the motif in peaks found in the entire aggregated single cell dataset^11^. We considered cells with a minimum of 2000 fragments per cell and a minimum of 10% of total fragments in peaks. To avoid biases from recovery of different numbers of cells for each sgRNA, we subsampled all sgRNA cell populations to 12 cells (the lowest number of cells for a single sgRNA in our K562 dataset), calculated the deviation *Z*-scores, and repeated this resampling process 1000 times to obtain deviation *Z*-scores for each sgRNA.

### Gene ontology analysis of differential EZH2 chromatin accessibility sites

In order to identify and annotate genomic regions that are differentially accessible in cells with *EZH2-targeting* sgRNAs, we aggregated equal numbers of single cells (*n* = 170 cells per sgRNA) for each of the three *EZH2* and non-targeting sgRNAs. We next binned the genome into 150 nt regions and identified all bins covered by all three *EZH2* sgRNAs and not covered by any of the three non-targeting sgRNAs. These bins were then mapped to the transcription start site of the closest genes. We used this (unranked) gene list (*n* = 3,740) as input for Gene Ontology enrichment analysis, with all human genes as a background set^44^.

### Differential accessibility at HOX loci and gene expression

To measure accessibility at HOX loci, *EZH2*-targeted and non-targeting single cells were downsampled to 100 cells, aggregated and fragments overlapping the *HOXA-D* loci were counted. Empirical *p*-values were calculated over 1000 bootstrap iterations. To select HOX genes for expression profiling, we compared CRISPR-sciATAC coverage in *EZH2* KO cells and NT cells. We computed the number of reads in each *HOX* gene body (including 500 nt flanking sequence on each side). We then selected the top 5 HOX genes with the most significant change in CRISPR-sciATAC coverage (Student’s *t*-test). Gene expression of HOX genes (*HOXA3, HOXA5, HOXA11, HOXA13, HOXD9*) and EZH2 following *EZH2* knock-out was quantified using quantitative qRT-PCR. Briefly, 1 million K562-Cas9 cells were infected with *EZH2* sgRNA 1-3 or NT sgRNA 1-3 at an MOI of ~0.1 for each of the 6 sgRNAs and grown in 6-well plates. At 24h post-infection, cells were selected in 1 μg/ml puromycin. Cells were harvested 10 days after transduction and lysed using TRIzol (Life Technologies), RNA was purified using Direct-zol (Zymo Research). We reverse-transcribed 1 μg of total RNA using random hexamer primers and RevertAid Reverse Transcriptase (Thermo Fisher) at 25°C for 10 min, 37°C for 60 min, and 95°C for 5 min. After cDNA synthesis qPCR reactions were performed using Luna Universal Probe qPCR Master Mix (NEB), custom primers and probes (IDT) were designed to detect each target gene and normalized to β-actin (ACTB) (see **Supplementary Table 2** for primer and probe sequences). All qPCRs were thermocycled on a ViiA 7 Real-Time PCR System (Applied Biosystems) as follows: initial denaturation at 95 °C for 1 min, then 40 cycles at 95 °C for 15 s, 60 °C for 30 s. Quantification was performed via the ΔΔC_t_ using 3 biological replicates and 4 technical (qPCR) replicates for each biological replicates.

### Identification of disrupted TF co-regulation in EZH2-targeted cells

To study disruption of co-regulation of TFs using CRISPR-sciATAC data, we implemented an approach similar to the one presented in Perturb-ATAC^27^. We compared correlations between differential accessibility patterns in each pair of TFs in cells that received non-targeting sgRNAs and cells that received EZH2 sgRNA. To ensure a comparison between single cells that received a successful perturbation, we considered a subset of EZH2-perturbed single cells that received the most effective sgRNA out of the three EZH2-trgeting sgRNAs (EZH2 sgRNA-3, see **Fig. 2g**), and from those cells we focused only on ones that showed increased accessibility in EZH2 binding sites (146/170 single cells, 85%). Since EZH2 is a key component of the Polycomb Repressive Complex 2 (PRC2) which generates heterochromatin where it binds, higher accessibility in EZH2 binding sites would be expected in EZH2-targeted cells due to failure in PRC function, as demonstrated in **Fig. 2c**. We calculate the Pearson correlation coefficient between each pair of 115 TFs (excluding binding sites of EZH2 from the analysis), over cells that received a non-targeting sgRNA and cells that received the EZH2 sgRNA. To identify changes in the correlation structure following EZH2 knock-out, we subtracted the coefficient calculated using single cells that received a non-targeting sgRNA from the coefficient calculated with EZH2-perturbed single cells.

### eQTL enrichment

To test if targeting chromatin modifiers resulted in changes in accessibility at SNPs associated with regulatory function through expression quantitative trait locus (eQTL) association testing, we utilized *cis*-eQTLs (SNP-gene combinations within 1 Mbp) from the eQTLGen consortium. The consortium performed association testing for 19,960 genes expressed in blood in 31,684 samples^45^. We considered the fraction of fragments in each single cell that overlap *cis*-eQTLs and compared these fractions for each population of single cells that received sgRNAs targeting a gene to the fractions in non-targeting cells using a Wilcoxon signed-rank test followed by a Benjamini-Hochberg multiple hypothesis correction. To identify specific *cis*-eQTLs with altered accessibility, we downsampled KDM6A single cells and non-targeting single cells to the same amount of cells (*n* = 737 cells) and focused on a subset of 7829 highly covered (≥ 50 reads) *cis*-eQTLs in the two cell populations combined. For each of these 7829 *cis*-eQTLs we considered the proportion of cells with a read covering the *cis*-eQTL in the KDM6A cell population (*n* = 921 cells) and in the non-targeting cell population (*n* = 737 cells) and performed a χ^2^ test of proportion. Allelic effects are from the Genotype-Tissue Expression (GTEx) database^46^. For each *cis*-eQTL, we show allele specific expression for the closest gene in whole blood samples. In cases where allele specific expression was not available in whole blood samples, we show the most significant association.

### Differential accessibility in enhancers and promoters

For each single cell, we calculated the fraction of reads that intersect with promoters and with enhancers, as defined by ENCODE (`wgEncodeAwgSegmentationCombinedK562.bed`, http://hgdownload.cse.ucsc.edu/goldenpath/hg19/encodeDCC/wgEncodeAwgSegmentation/).

We then compared the fractions in each gene knock-out cell population with the fractions in the non-targeting cell population. To find significant differences in reads in promoters/enhancers (versus non-targeting), we performed a two-sample Wilcoxon test and the *p*-values were adjusted for multiple hypothesis testing using a Benjamini-Hochberg false-discovery rate (*q* ≤ 0.1).

### Nucleosome dynamics at TFBS, promoters and enhancers

To investigate nucleosome dynamics around TFBSs, we first subset the ATAC fragments into fragments putatively spanning one nucleosome (mono-nucleosome fragments, 147 – 280 bp^36^). We next calculated coverage profiles around TFBSs sites with `BEDTools`^47^. We focused on TFBSs that had two nucleosomes spanning them symmetrically as seen in our data. The TFBS selected for this analysis thus reflect those whose nucleosome positions are strongly bimodal. We chose these sites by calculating an nucleosome-free region (NFR) score for each, a metric to assess bi-modality^48^. Specifically, we take the difference in average base-pair coverage between flanking regions (50 to 150 bp upstream and downstream of site) and the central region (50 bp across site center). We focused on a small subset of 7 TFs with NFR score of less than zero, and indeed, for the binding sites of TFs, we observe strong bimodality. These TFs have also been previously shown to have a bimodal profile^21^.

ATAC-seq fragment coverage plots were smoothed using the `smooth.spline` function (from the R package `stats`), with smoothing parameter `spar` = 0.8. Next, positions of maximum coverage upstream and downstream of motif centers were used to estimate nucleosome location. To determine expansion, we first calculated the distance between the upstream and downstream nucleosomes at a particular TFBS. Then, this distance was compared to non-targeting cells, to obtain a positive (expansion) or negative (compaction) score. Empirical *p*-values for each score were generated using a label-permutation test, where non-targeting and knock-out labels were randomly shuffled while keeping group size constant to avoid biases from different numbers of cells in the non-targeting and knock-out cell populations. Labels were shuffled 10,000 times and in each iteration the distance between the upstream and downstream nucleosomes was measured, to create a null distribution to which the true distance was compared.

For each TF, we calculate mononucleosomal coverage profiles separately for sites located in promoters or enhancers, as defined by UCSC (`wgEncodeAwgSegmentationCombinedK562.bed`, http://hgdownload.cse.ucsc.edu/goldenpath/hg19/encodeDCC/wgEncodeAwgSegmentation/).

### Statistical analysis

Data between two groups were analyzed using a two-tailed unpaired *t*-test or a non-parametric Wilcoxon signed-rank test. The *p* values and statistical significance were estimated for all analyses. Corrections for multiple-hypothesis testing was performed using the Benjamini-Hochberg approach^49^. In all the box plots, the central rectangle in the plot covers the first to the third quartile (the interquartile range, or IQR) and the bold line is the median. The whiskers are defined as: *whisker_upper_* = *min*(*max*(*x*), *Q_3_* + 1.5 × *IQR*) and *whisker_lower_* = *max(min(x), Q_1_* −1.5 × *IQR*). All statistical analyses were performed in R/RStudio.

### Data and resource availability

Processed and raw data can be downloaded from NCBI GEO (PRJNA674902, GSE161002).

