## Supplementary Figures and Sequences for "Scalable pooled CRISPR screens with single-cell chromatin accessibility profiling"

**Supplementary tables**

| | |
|---|---|
| Supplementary Table 1. | Number of cells in each step of the CRISPR-sciATAC protocol. |
| Supplementary Table 2. | Sequences of oligonucleotides for CRISPR-sciATAC, CRISPR libraries and qRT-PCR. |
| Supplementary Table 3. | Gene and sgRNA enrichment from essentiality screen. |
| Supplementary Table 4. | ENCODE ChIP data sources. |
| Supplementary Table 5. | Histone mark differential accessibility. |
| Supplementary Table 6. | GO enrichment results of differential accessibility in *EZH2*-targeted cells. |
| Supplementary Table 7. | Transcription factor binding site differential accessibility. |
| Supplementary Table 8. | Cost comparison between CRISPR-sciATAC and Perturb-ATAC protocols. |
| Supplementary Table 9. | Time comparison between CRISPR-sciATAC and Perturb-ATAC protocols. |

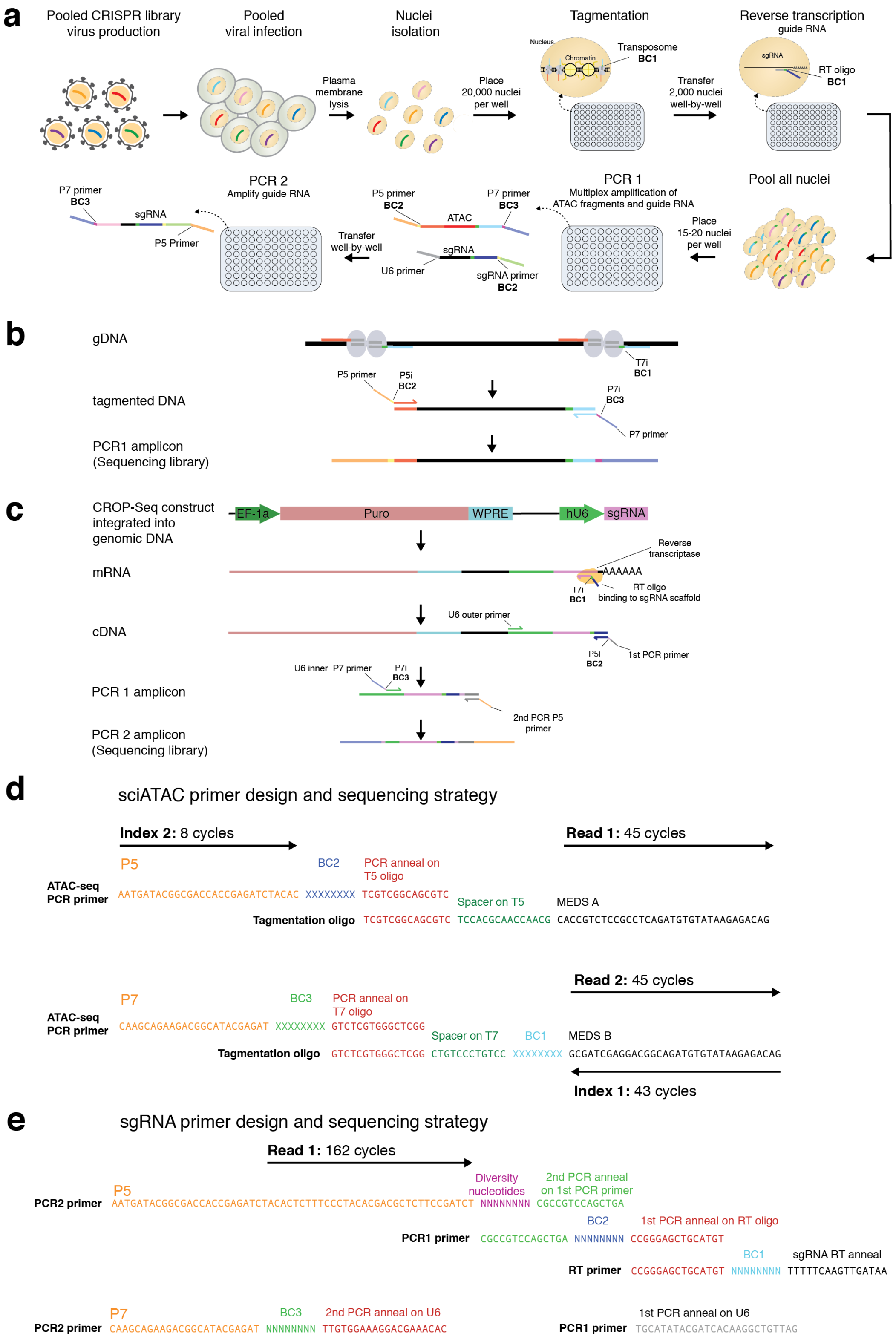


**Supplementary Figure 1. CRISPR-sciATAC library preparation and sequencing. (a)** Workflow for CRISPR-sciATAC. BC, barcode. Cell barcodes consist of a unique combination of BC1, BC2, and BC3. (**b**) CRISPR-sciATAC schematic for ATAC-seq library preparation. (**c**)

CRISPR-sciATAC schematic for sgRNA library preparation. (**d**) CRISPR-sciATAC primer design and sequencing strategy for ATAC fragments. (**e**) CRISPR-sciATAC primer design and library sequencing strategy for sgRNA amplicons. Staggered P5 oligos were introduced in the library preparation to introduce sequence diversity. BC 1, 2, and 3 are matched for ATAC-seq and sgRNA libraries, e.g. the ATAC-seq BC 1 in well A1 in the 96-well plate where tagmentation is performed is the same as the sgRNA BC 1 in well A1 in the 96-well plate where reverse transcription is performed.

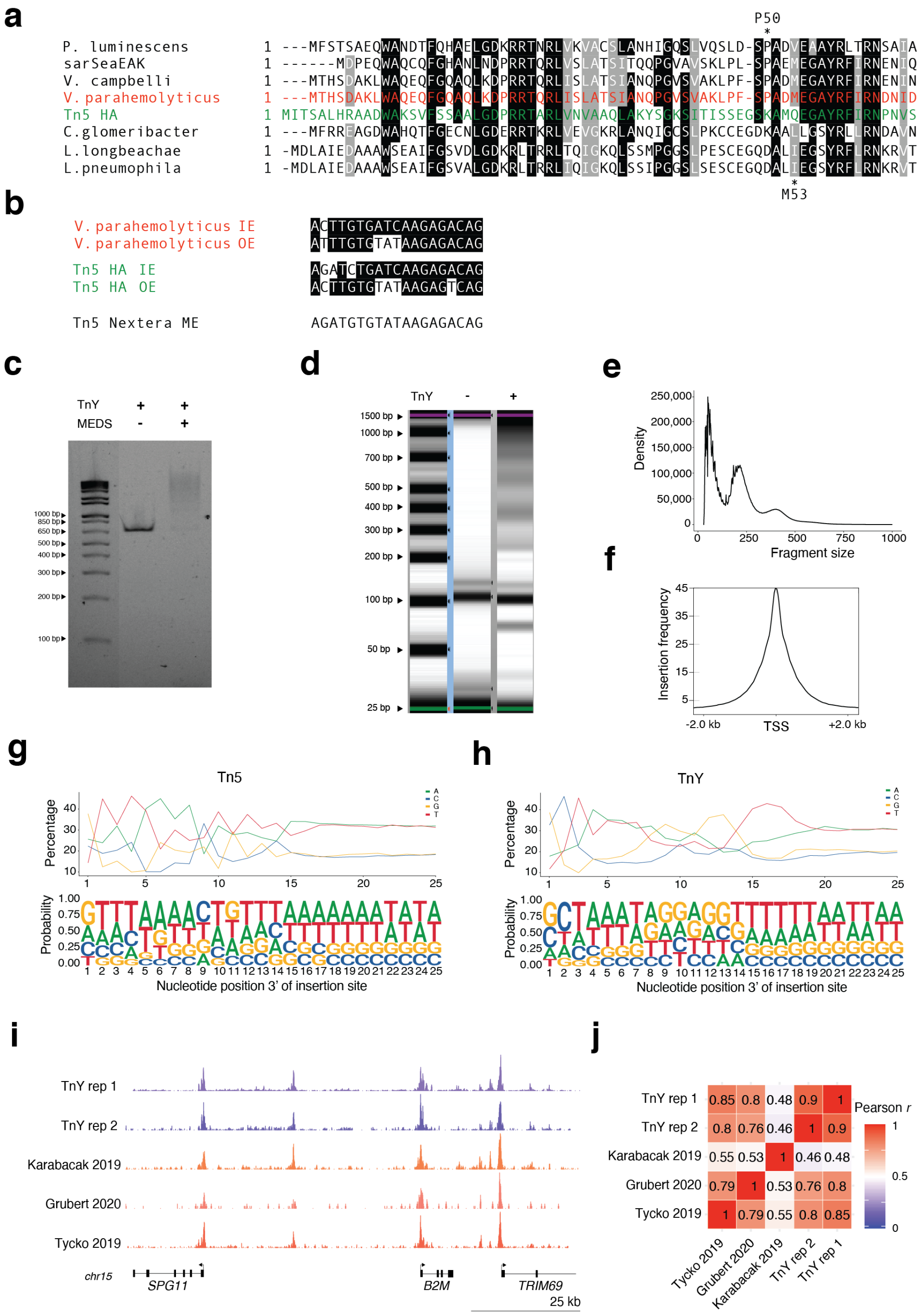


**Supplementary Figure 2. Comparison of TnY and Tn5 transposases.** (**a**) Alignment results of various bacterial transposases with a high-activity variant of Tn5 (Tn5_HA). Shared amino acids are shaded in black. Amino acids with similar properties are shaded in grey. Multiple alignment was done with ClustalW[1]. (**b**) Alignment of *V. parahemolyticus* transposon end sequences to those of the Tn5 transposon. Tn5 Nextera mosaic end (ME) sequence is also depicted. IE, inside end. OE, outside end. (**c**) Migration of ~700 bp PCR product after incubation with unloaded TnY or with TnY loaded with MEDS. (**d**) Capillary electropherogram of bulk tagmentation of K562 cells using TnY and a no-transposase negative control. The characteristic nucleosomal banding pattern is only visible in the condition with transposase. (**e**) Fragment size distribution and (**f**) ATAC-seq fragments insertions at transcription start sites (TSS) obtained from bulk tagmentation of K562 cells using TnY. (**g-h**) Nucleotide frequency plot (*upper panel*) and DNA sequence logo (*lower panel*) showing insertion bias of Tn5 (**g**) and TnY (**h**). (**i**) Comparison of a TnY bulk ATAC-seq dataset from K562 cells and three previously published K562 Tn5 ATAC-seq datasets (Karabacak 2019: GSM2902635[2], Grubert 2020: GSM4130894[3], Tycko 2019: GSM3770765[4]) at a specific genomic locus (**j**) Correlation between ATAC accessibility averaged over 10 kb genomic bins for the datasets shown in (**i**).

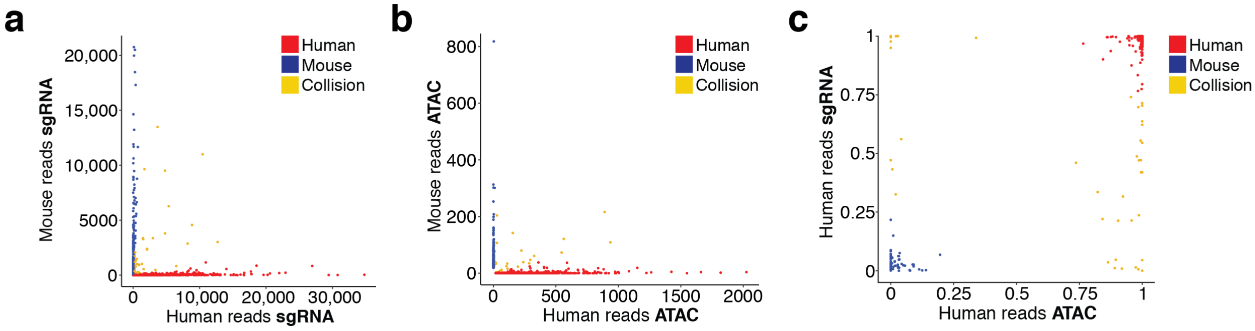


**Supplementary Figure 3. Separation of human and mouse single-cell ATAC and sgRNA reads in the species-mixing experiment.**

(**a**) Guide RNA reads mapping to human or mouse CRISPR libraries (*n* = 1986 cells). (**b**) ATAC reads mapping to human or mouse genomes (*n* = 721 cells). In this plot, a single cell that had >10-fold the number of reads of the average read number over all cells removed for display purposes. (**c**) Concordance between ATAC and sgRNA reads mapping to the human and mouse genomes and human and mouse sgRNA libraries for each cell (*n* = 496 cells).

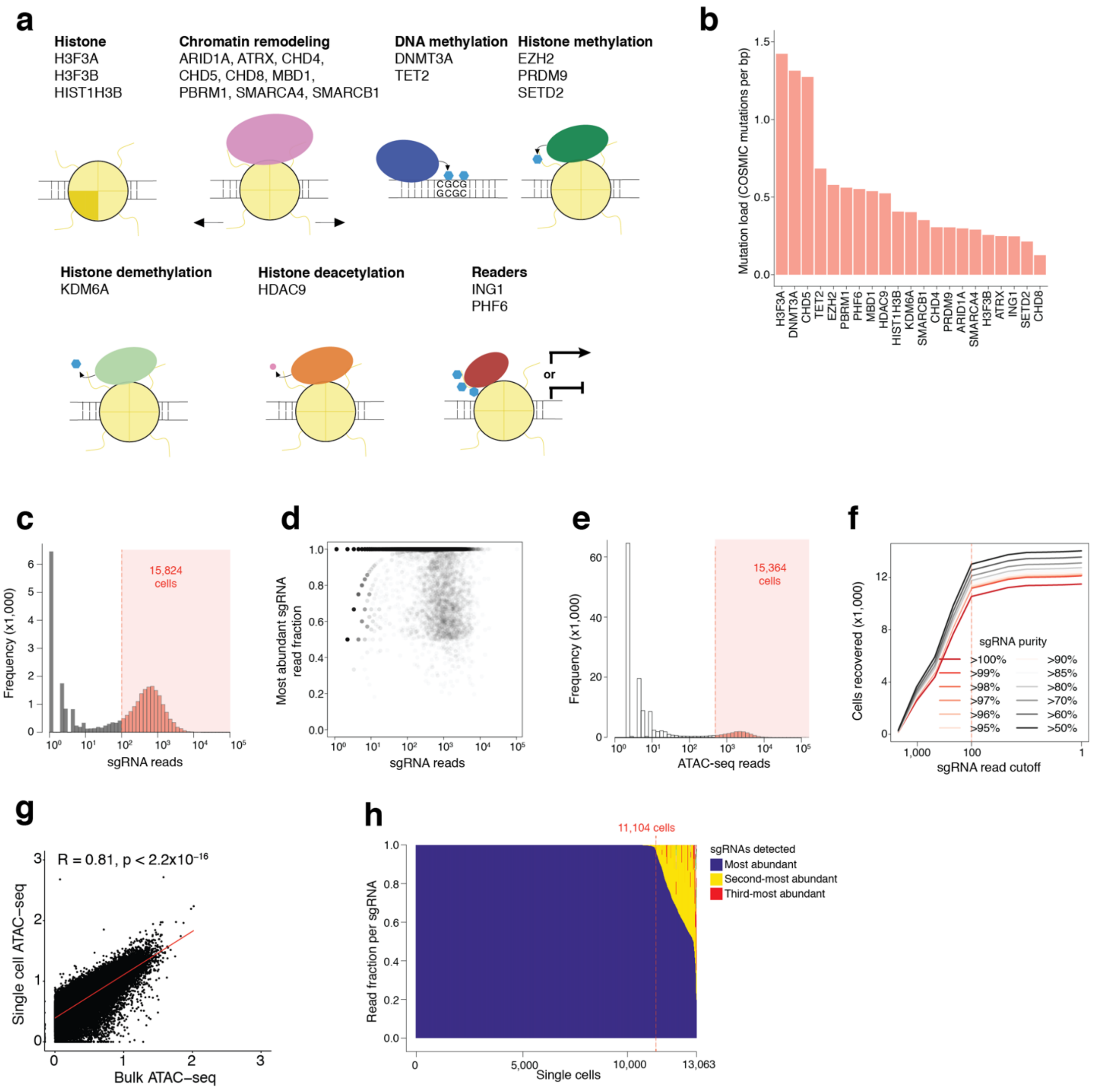


**Supplementary Figure 4. A pooled screen of 21 commonly mutated chromatin modifiers using CRISPR-sciATAC.** (**a**) Chromatin modifiers targeted in the 21 gene CRISPR library. (**b**) Mutation load for genes targeted in the chromatin modifier CRISPR library. For each of the chromatin modifiers targeted in the CRISPR library, mutation load is calculated by dividing the number of exonic mutations (in the COSMIC database[5]) by the gene length. Selected genes represent the top 20 most frequently mutated chromatin modifiers, as defined by mutation load, plus *CHD8*. (**c**) sgRNA reads per cell. 15,824 cells had at least 100 sgRNA reads (55% of all unique barcode combinations). (**d**) Proportion of sgRNAs with the highest read count per cell compared to the number of total sgRNA reads per cell. (**e**) Unique ATAC-seq reads per cell. 15,364 cells had at least 500 unique reads (12% of all unique barcode combinations). (**f**) Comparison of number of filtered ATAC-seq cells (filtering for ≥500 unique ATAC-seq reads) with the number sgRNA reads across different sgRNA purity thresholds. (**g**) Correlation of normalized accessibility between bulk ATAC-seq and single-cell ATAC-seq (aggregated over 11,104 single cells with both ATAC-seq and sgRNA capture). The normalized accessibility is averaged over 10 kb genomic bins and a linear regression line is shown in red ($r_p = 0.81$, $p < 2.2\text{x}10^{-16}$). (**h**) Representation of sgRNAs within each single cell. The most abundant sgRNA within each cell is colored in blue, the second most abundant sgRNA within each cell is colored in yellow and the third most abundant sgRNA in each cell is colored in red. 11,104 cells with ≥99% sgRNA reads from a single sgRNA were chosen for further analyses.

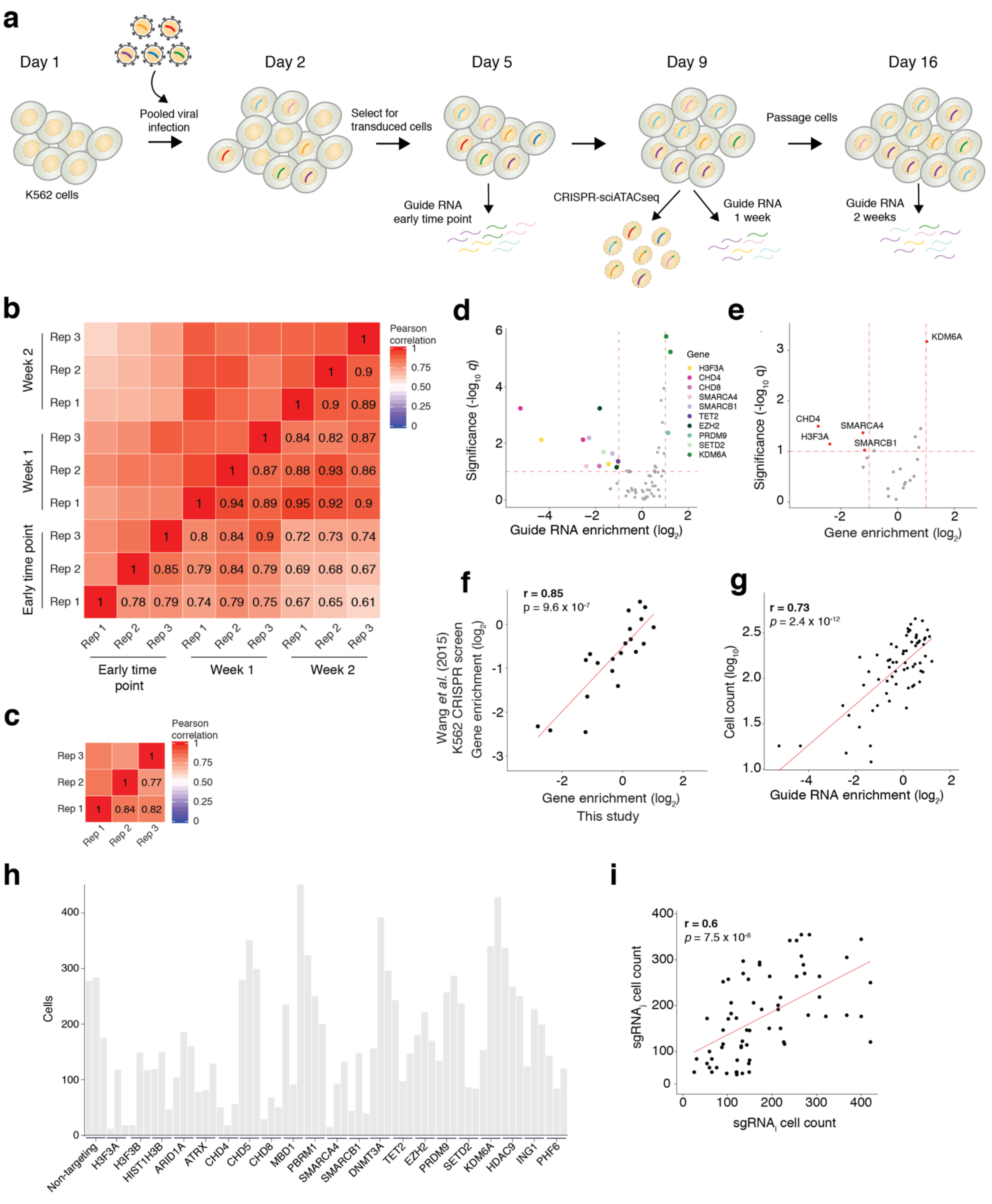


**Supplementary Figure 5. CRISPR pooled screen enrichment/dropout analysis.** (**a**) Screen timeline indicating both traditional CRISPR screen readout and CRISPR-sciATAC. (**b**) Pearson correlation between normalized read counts ($n$ = 3 transduction replicates for all samples). (**c**) Pearson correlation of the enrichment of library sgRNAs between Week 2 and Early Time Point samples in the 3 biological replicates. (**d**) Volcano plot of sgRNA-level enrichment (defined as

$log_2$ fold-change between week 2 and the early time point) and significance. sgRNAs highlighted in color have $|\log_2(\textit{sgRNA enrichment})| \geq 1$ and $q \leq 0.1$. Enrichment values are averaged over the three transduction replicates. Colors correspond to the gene function depicted in *a*. (**e**) Volcano plot of gene-level enrichment score and Bonferroni-corrected *p*-values ($-\log_{10} q$). The gene-level enrichment is computed as the average enrichment over biological replicates and then over sgRNAs for each gene. Genes highlighted in red have $|\log_2(\textit{gene-level enrichment})| \geq 1$ and $q \leq 0.1$. (**f**) Correlation of gene-level enrichment between this study and a previous genome-scale CRISPR screen in K562 cells[6]. (**g**) Scatterplot of sgRNA enrichment from the traditional CRISPR screen readout and cell counts (single cell barcodes) in the CRISPR-sciATAC screen. (**h**) Single cells per sgRNA from the CRISPR-sciATAC experiment in K562 cells. (**i**) Correlation between cell counts (single cell barcodes) for every pair of sgRNAs targeting the same gene.

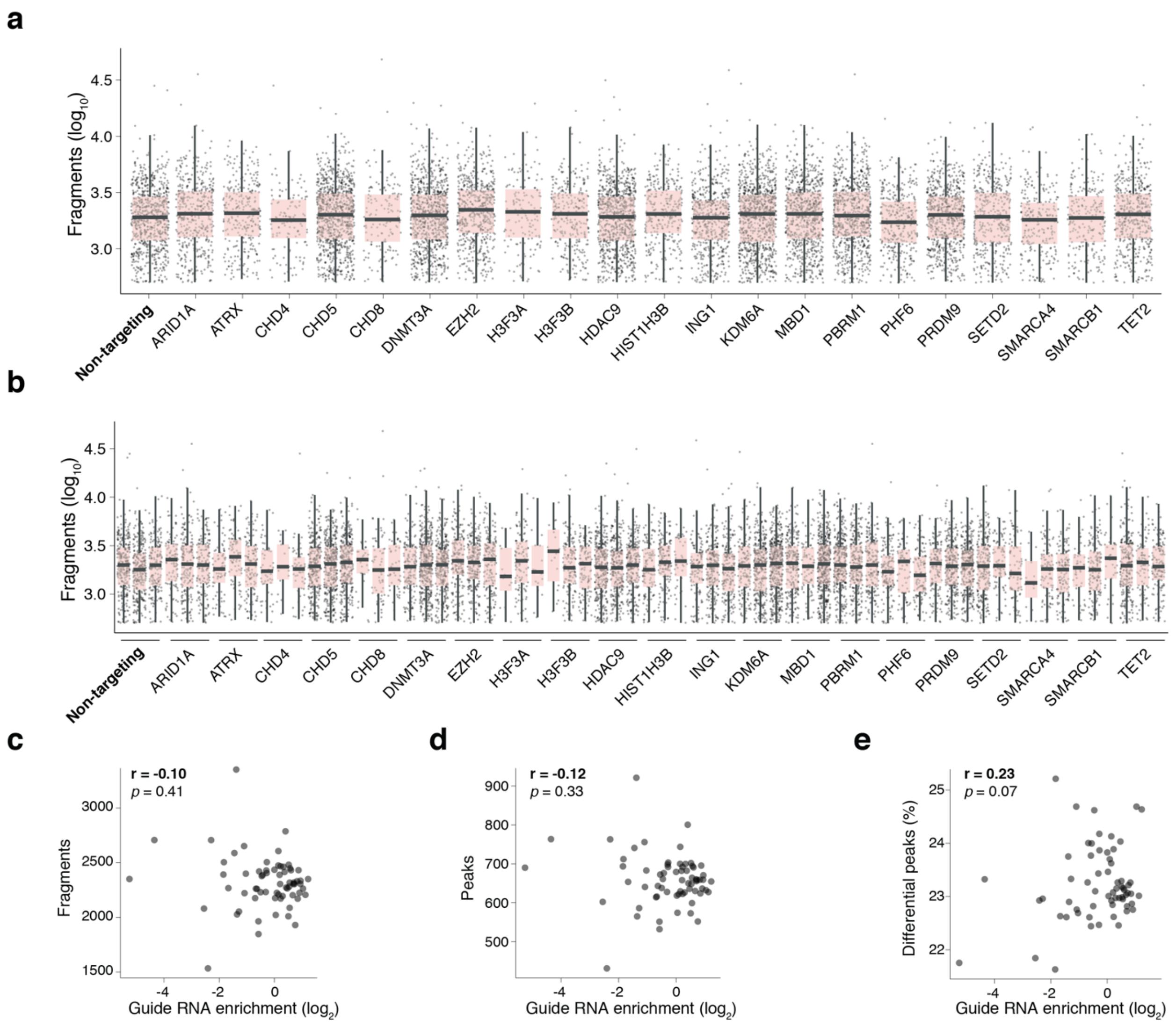


**Supplementary Figure 6. ATAC-seq fragments counts by sgRNA and target gene.** Number of ATAC-seq fragments in each targeted gene (**a**) and sgRNA (**b**) cell population. Each dot represents a single cell. The number of fragments from cells of each sgRNA were compared to the number of fragments in non-targeting (NT) cells. There were no significant changes in fragment counts observed (Wilcoxon rank-sum test, $p \leq 0.1$ following a Bonferroni correction). (**c**) Scatterplot of ATAC-seq fragments per sgRNA (averaged over cells) and sgRNA enrichment. (**d**) Scatterplot of peaks called per sgRNA (averaged over cells) and sgRNA enrichment. (**e**) Scatterplot of the percent of differential peaks per sgRNA and sgRNA enrichment. The fraction of differential peaks is defined as the proportion of peaks that exist only in cells that received that sgRNA and are not found in cells that receive NT sgRNAs. For panels *c*, *d* and *e*, all correlations shown are Pearson correlations and none are significant.

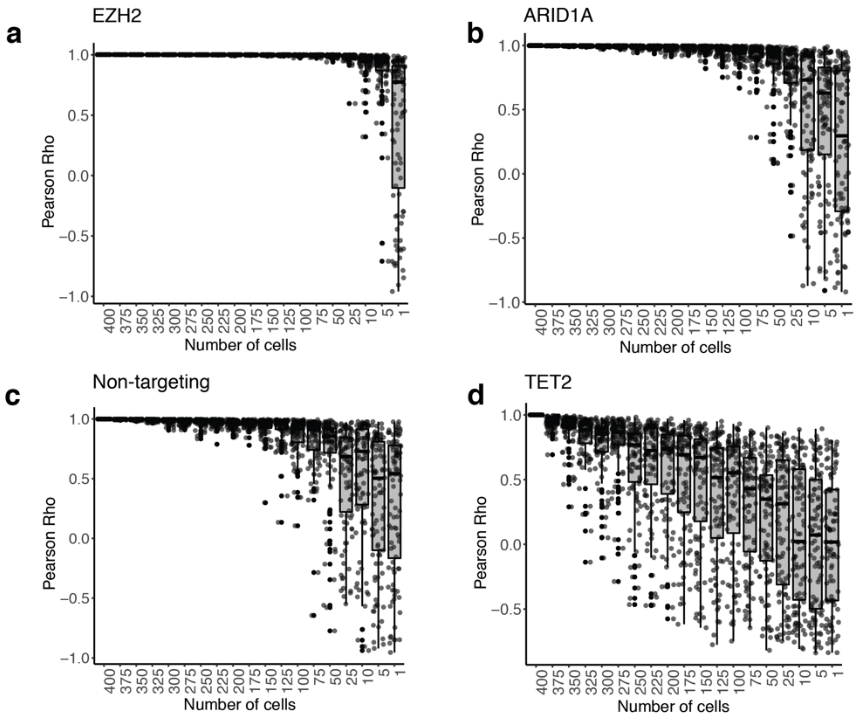


**Supplementary Figure 7. Correlation of downsampled cell populations with the aggregated (pseudo-bulk) dataset.** Pearson correlation between averaged histone mark *Z*-score profiles of the indicated number of single cells and the average profile of 400 single cells that received the same perturbation. For each cell number, we randomly sampled cells (from a total of 400 single cells) without replacement. We performed this resampling 200 times for each cell number. Data is shown for cells transduced with *EZH2*-targeting (**a**), *ARID1A*-targeting (**b**), non-targeting (**c**) and *TET2*-targeting sgRNAs (**d**).

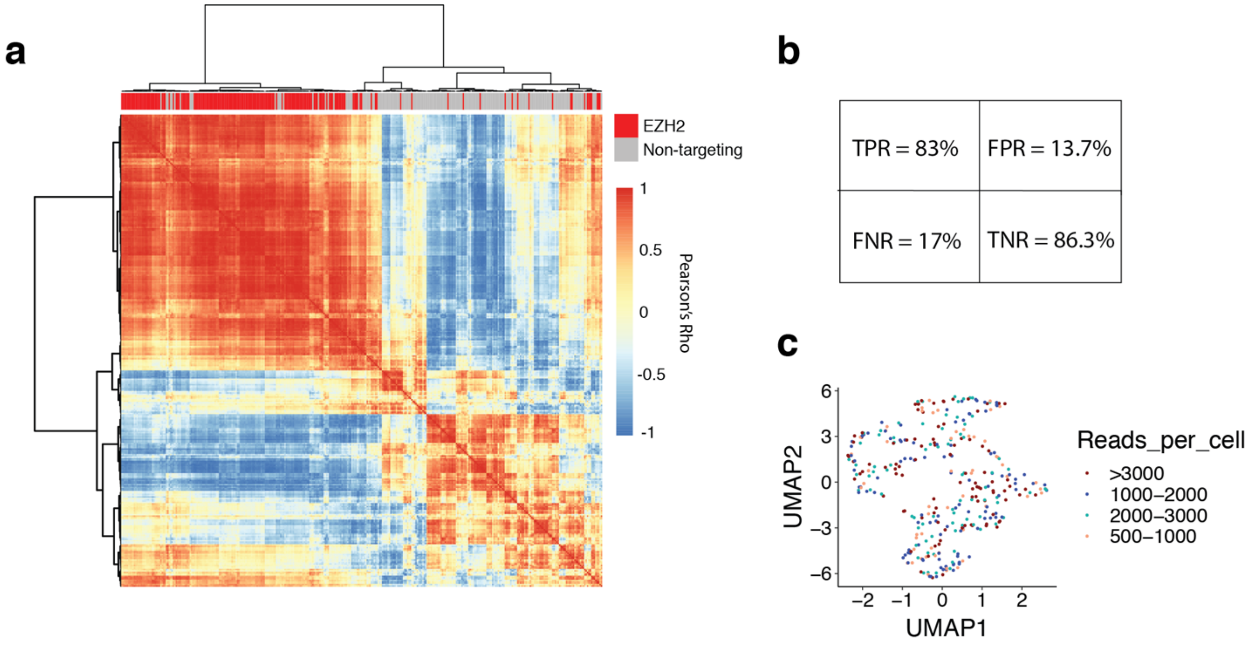


**Supplementary Figure 8. Clustering of single cells with either EZH2-targeting or non-targeting sgRNAs.** (**a**) Hierarchical clustering of single cells transduced with either *EZH2*-targeting or non-targeting sgRNAs. (**b**) Confusion matrix showing True Positive Rate (TPR), False Positive Rate (FPR), False Negative Rate (FNR) and True Negative Rate (TNR) for the clustering presented in panel *a* when cutting the dendrogram at $k = 2$. (**c**) The same UMAP representation as shown in **Fig. 2d** with the cells colored by the number of reads per cell.

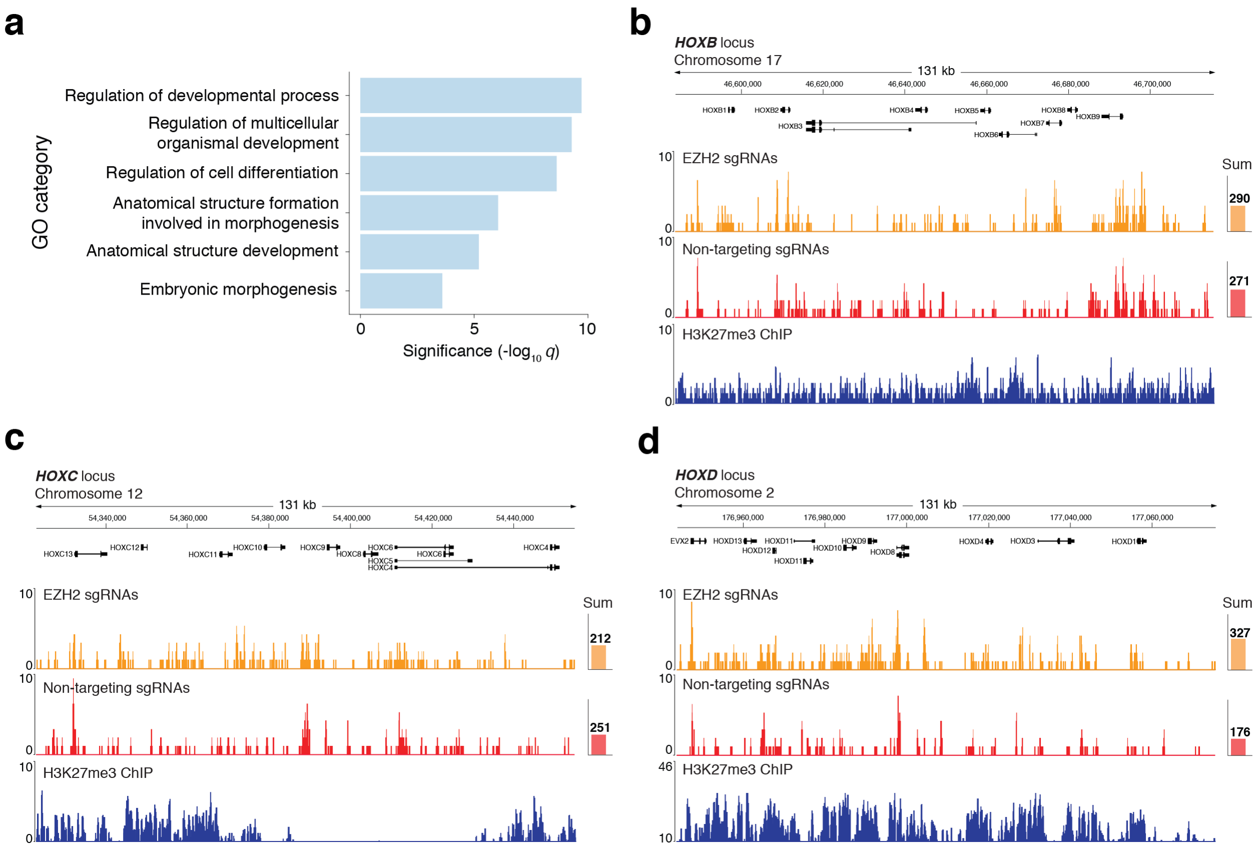


**Supplementary Figure 9. ATAC-seq fragments at HOX genes in cells with EZH2-targeting or non-targeting sgRNAs**. (**a**) Gene ontology (GO) terms enriched for genes close to genomic regions with differential accessibility following *EZH2* disruption. Shown are selected GO terms with significant enrichment; the full list is given in **Supplementary Table 6**. (**b**) - (**d**) CRISPR-sciATAC fragments mapping to the *HOXB* (**b**), *HOXC* (**c**), and *HOXD* (**d**) loci in cells transduced with *EZH2*-targeting (*orange*) and non-targeting (*red*) sgRNAs ($n = 510$ cells per condition). K562 H3K27me3 ChIP-seq coverage is shown at the bottom (*blue*). The sum of all ATAC fragments in each locus in *EZH2*-targeted and non-targeting aggregated single cells is shown on the right.

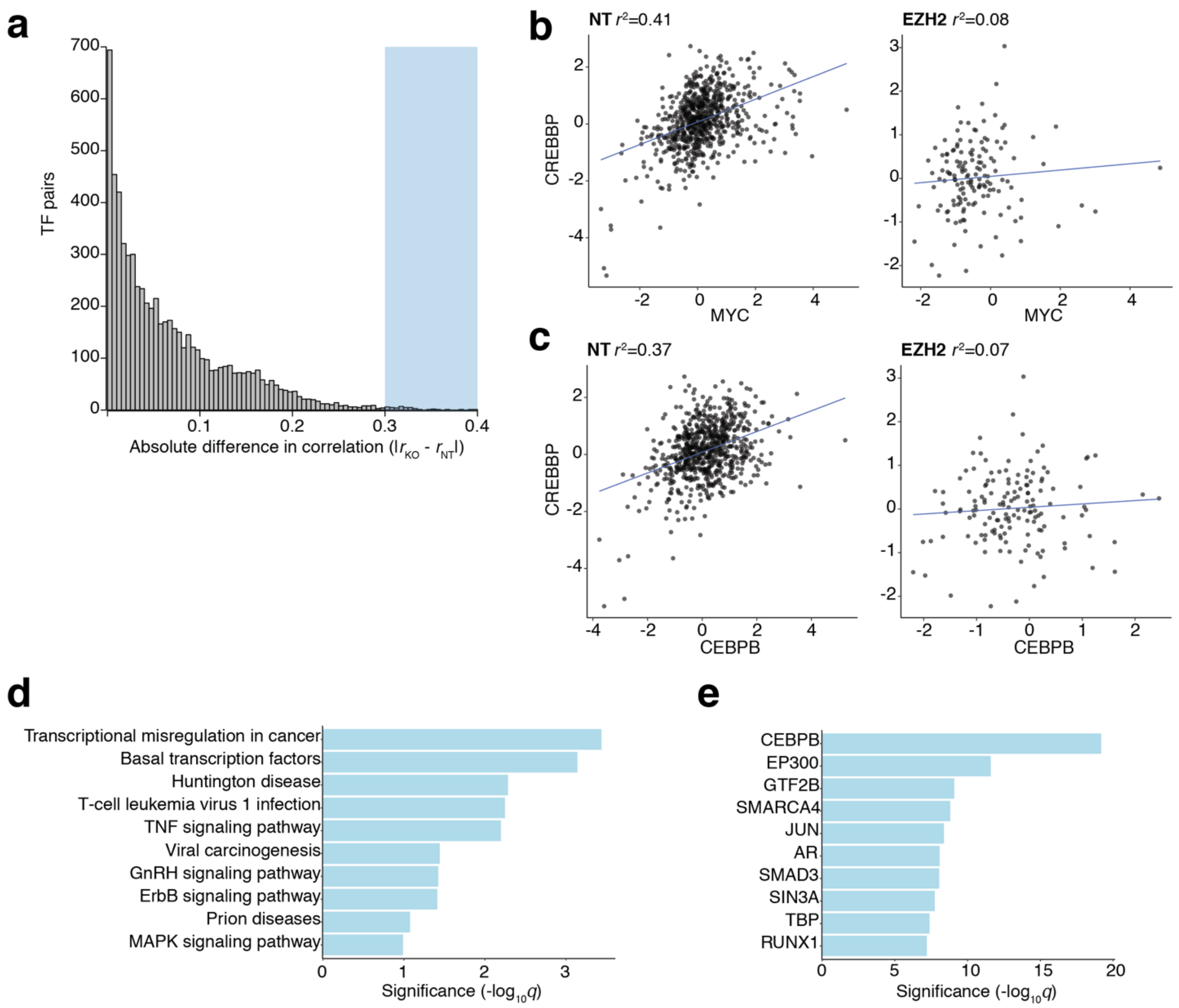


**Supplementary Figure 10. Disruption of CREBBP co-accessibility following EZH2 knock-out.** (**a**) Histogram of the absolute value of the difference in *TFBS correlation* between cells receiving EZH2-targeted and non-targeting sgRNAs, over 6,555 possible TF pairs. Here, we define the *TFBS correlation* as the Pearson correlation between single-cell chromatin accessibility *Z*-scores for all binding sites of 2 TFs across single cells (i.e. $TF_1$ vs. $TF_2$). (**b**) The *TFBS correlation* for CREBBP and MYC binding sites in cells that received a non-targeting sgRNA (*left*) or EZH2-targeting sgRNA (*right*). (**c**) The *TFBS correlation* for CREBBP and CEBPB binding sites in cells that received a non-targeting sgRNA (*left*) or EZH2-targeting sgRNA (*right*). (**d**) Most-enriched KEGG Pathways for TFs exhibiting a disruption in binding site co-accessibility with CREBBP binding sites (*n* = 40 transcription factors)[7]. (**e**) Most-enriched protein-protein interactions from Enrichr for TFs exhibiting a disruption in binding site co-accessibility with CREBBP binding sites (*n* = 40 transcription factors)[8].

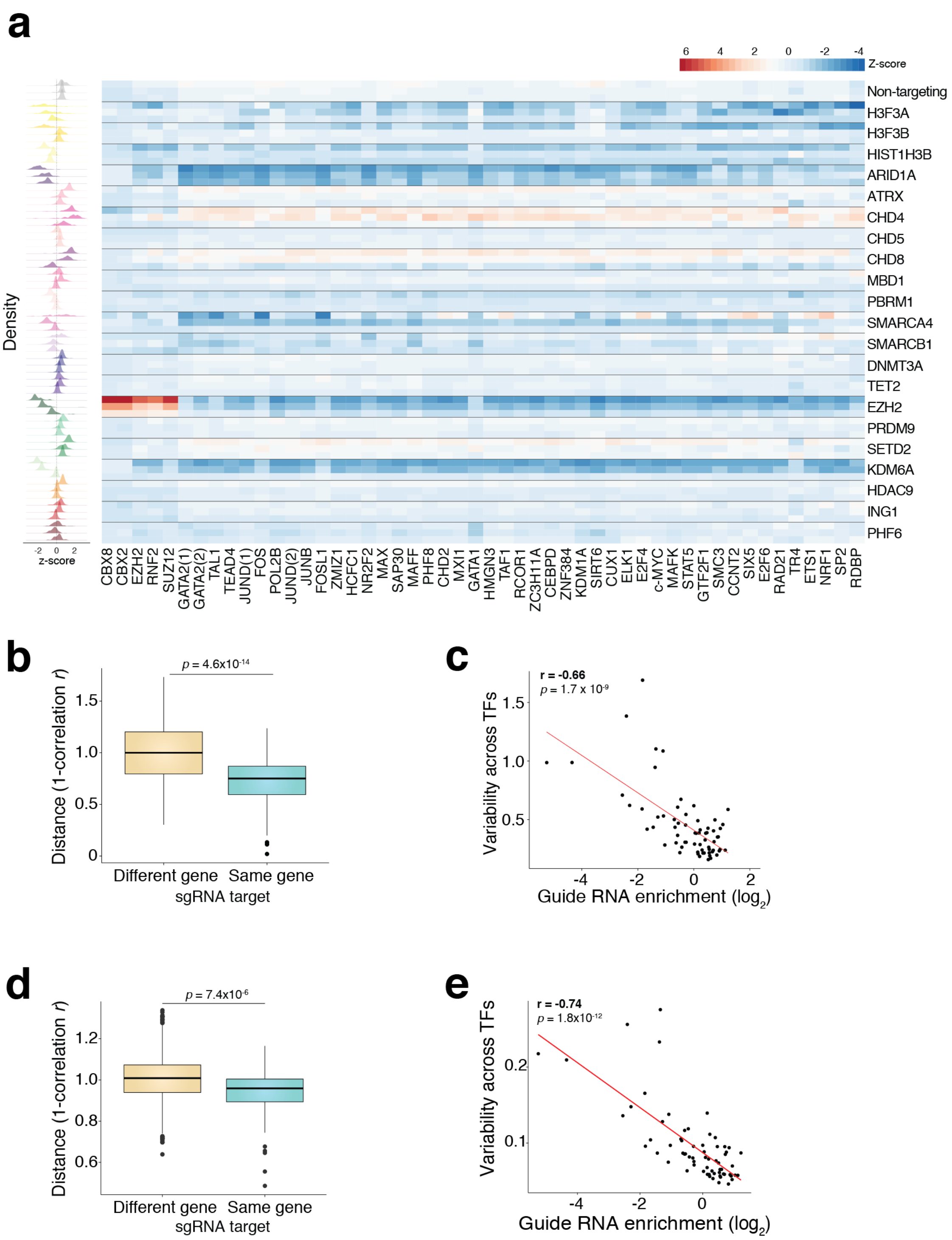


**Supplementary Figure 11. Differential accessibility in TF binding sites.** (**a**) Heatmap showing accessibility at transcription factor binding sites (TFBSs) for the different sgRNAs ($n$ = 3 sgRNA per gene). The fraction of reads in TFBSs was calculated for each single cell. Fractions are standardized for each TFBS over all single cells and then averaged over all single cells for each sgRNA cell population. Shown are the 50 transcription factors with the most significant differences in accessibility. The histograms on the left show the distribution of *Z*-scores for each sgRNA; colors correspond to the gene function depicted in **Supplementary Fig. 4a**. The *Z*-scores are computed over all cells in the screen. (**b**) Distances in the TFBS accessibility profiles shown in panel *a* between sgRNAs targeting different genes and sgRNAs targeting the same gene. The distance metric used is 1-(Pearson correlation). (**c**) Scatterplot of guide-level enrichment from the pooled screen readout and the standard deviation (across sgRNAs) of TFBS accessibility profiles shown in panel *a*. (**d**) Distances in the TFBS accessibility profiles, calculated using the `chromVAR` package[9], between sgRNAs targeting different genes and sgRNAs targeting the same gene. The distance metric used is 1-(Pearson correlation). (**e**) Scatterplot of guide-level enrichment from the pooled screen readout and the standard deviation (across sgRNAs) of `chromVAR` TFBS accessibility profiles.

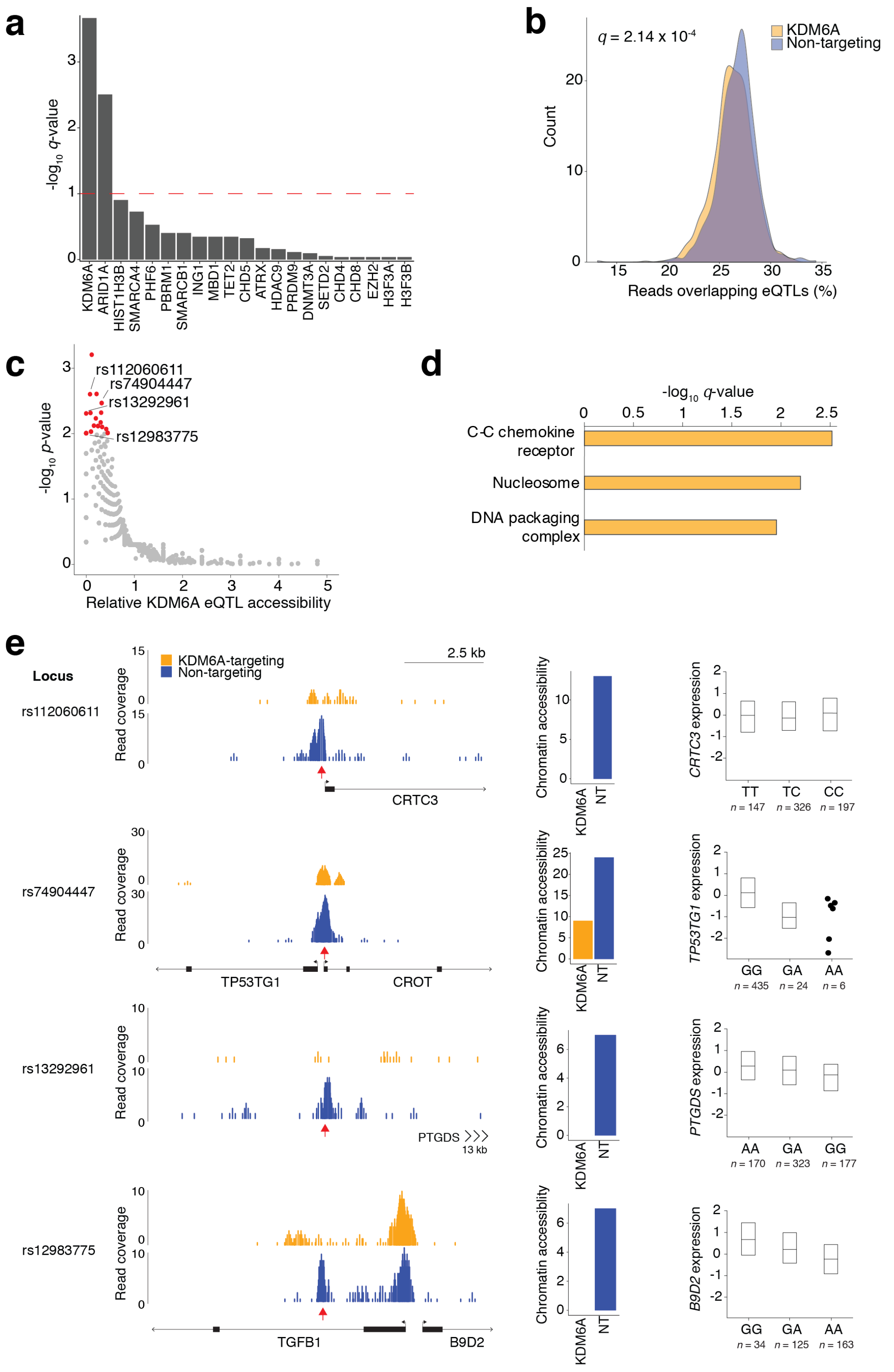


**Supplementary Figure 12. Changes in chromatin accessibility at blood *cis*-eQTLs. (a)** Significance of the change in percent of reads (ATAC fragments) overlapping blood *cis*-eQTLs in each of the knock-out cell populations in comparison to the non-targeting cell population. We tested for significant differences using Student's *t* test (two-tailed) and corrected for multiple comparisons using the Benjamini-Hochberg false-discovery rate (FDR) correction. The dashed red line represents a FDR significance cutoff with *q*-value = 0.1. (**b**) The percent of reads (ATAC fragments) overlapping at least one blood *cis*-eQTL in KDM6A-targeted cells. Compared to cells that receive a non-targeting sgRNA, KDM6A-targeted cells have reduced chromatin accessibility at blood *cis*-eQTLs. (**c**) Scatterplot of relative chromatin accessibility of KDM6A-targeted cells and significance ($\chi^2$ test of proportions) in a subset of blood *cis*-eQTLs with high coverage ($\geq 50$ reads) in either NT or KDM6A cells ($n = 7829$ *cis*-eQTLs). In this analysis, we define relative chromatin accessibility of KDM6A-targeted cells as the ratio of reads overlapping each site in the KDM6A-perturbed cell population and in the non-targeting cell population (normalized by the number of cells in each cell population). Red dots represent eQTLs which are differentially accessible in KDM6A-targeted cells with nominal significance. (**d**) Gene ontology (GO) terms enriched for genes (eGenes) whose expression is affected by differentially accessible cis-eQTLs. (**e**) Four differentially accessible eQTLs highlighted in panel *c*. *Left*: ATAC read pileups comparing accessibility between cells with a KDM6A-targeting or non-targeting sgRNA at select eQTLs (red arrows). *Center*: Number of fragments in eQTLs for cells with a KDM6A-targeting or non-targeting sgRNA. *Right*: For the eQTL, gene expression across different haplotypes from the Genotype-Tissue Expression (GTEx) dataset[10].

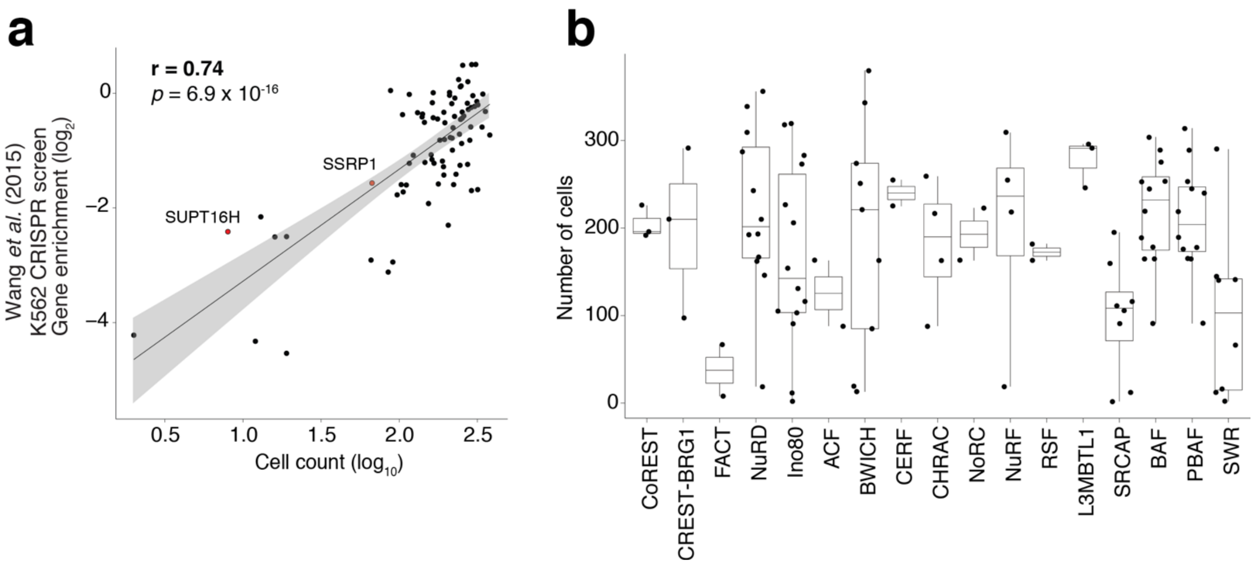


**Supplementary Figure 13. Chromatin remodeling complex essentiality.** (**a**) Correlation of single-cell barcodes obtained in the CRISPR-sciATAC screen and gene enrichment scores from a previous genome-scale CRISPR screen in K562 cells[6]. FACT subunits SUPT16H and SSRP1 are labeled with points colored in red. (**b**) Boxplot of the number of single-cell barcodes obtained per gene for each chromatin remodeling complex. Each dot represents a subunit in the complex as defined by EpiFactors[11].

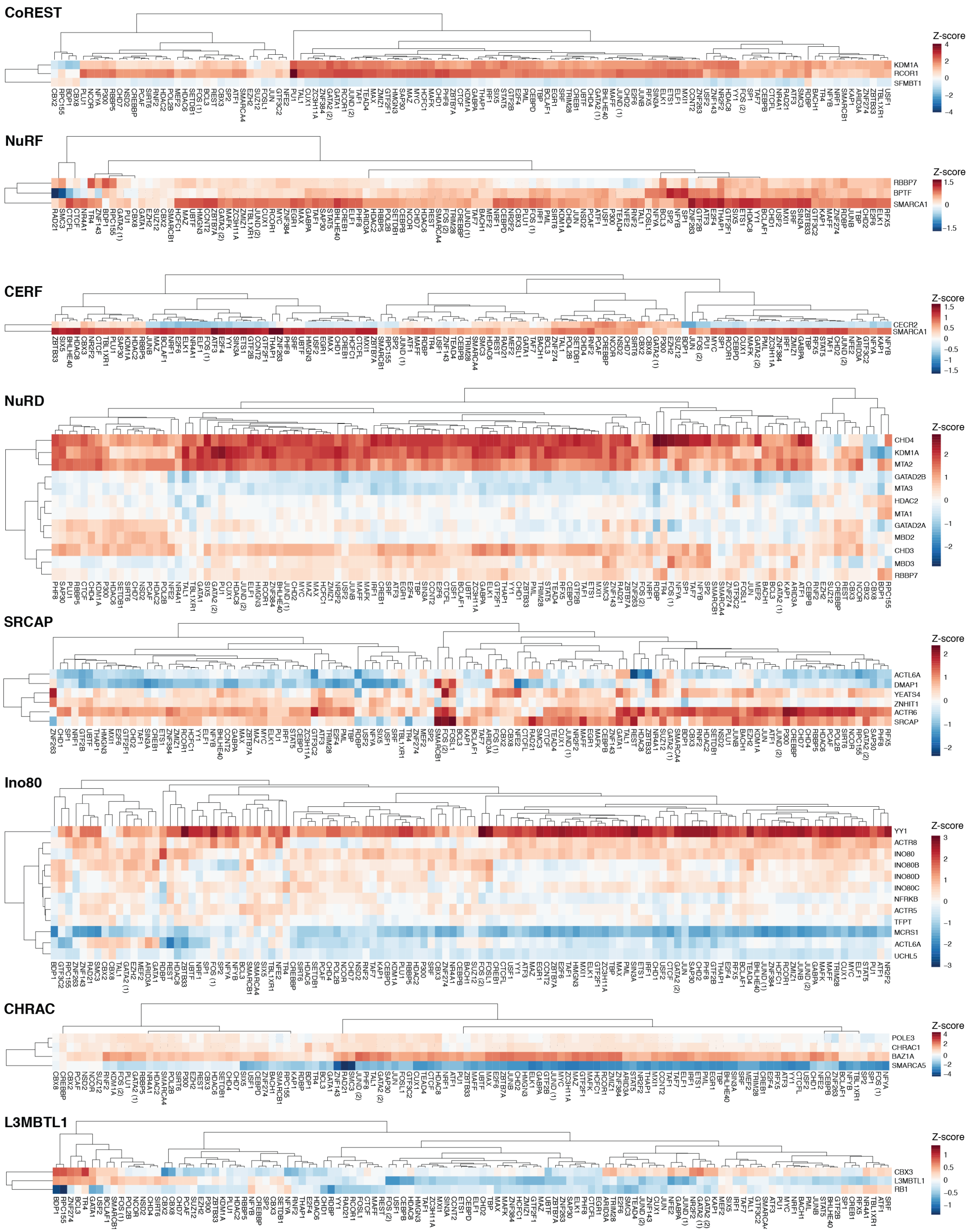

CoREST
KDM1A
RCOR1
SFMBT1
Z-score
NuRF
RBBP7
BPTF
SMARCA1
Z-score
CERF
CECR2
SMARCA1
Z-score
NuRD
CHD4
KDM1A
MTA2
GATAD2B
MTA3
HDAC2
MTA1
GATAD2A
MBD2
CHD3
MBD3
RBBP7
Z-score
SRCAP
ACTL6A
DMAP1
YEATS4
ZNHIT1
ACTR6
SRCAP
Z-score
Ino80
YY1
ACTR8
INO80
INO80B
INO80D
INO80C
NFRKB
ACTR5
TFPT
MCRS1
ACTL6A
UCHL5
Z-score
CHRAC
POLE3
CHRAC1
BAZ1A
SMARCA5
Z-score
L3MBTL1
CBX3
L3MBTL1
RB1
Z-score

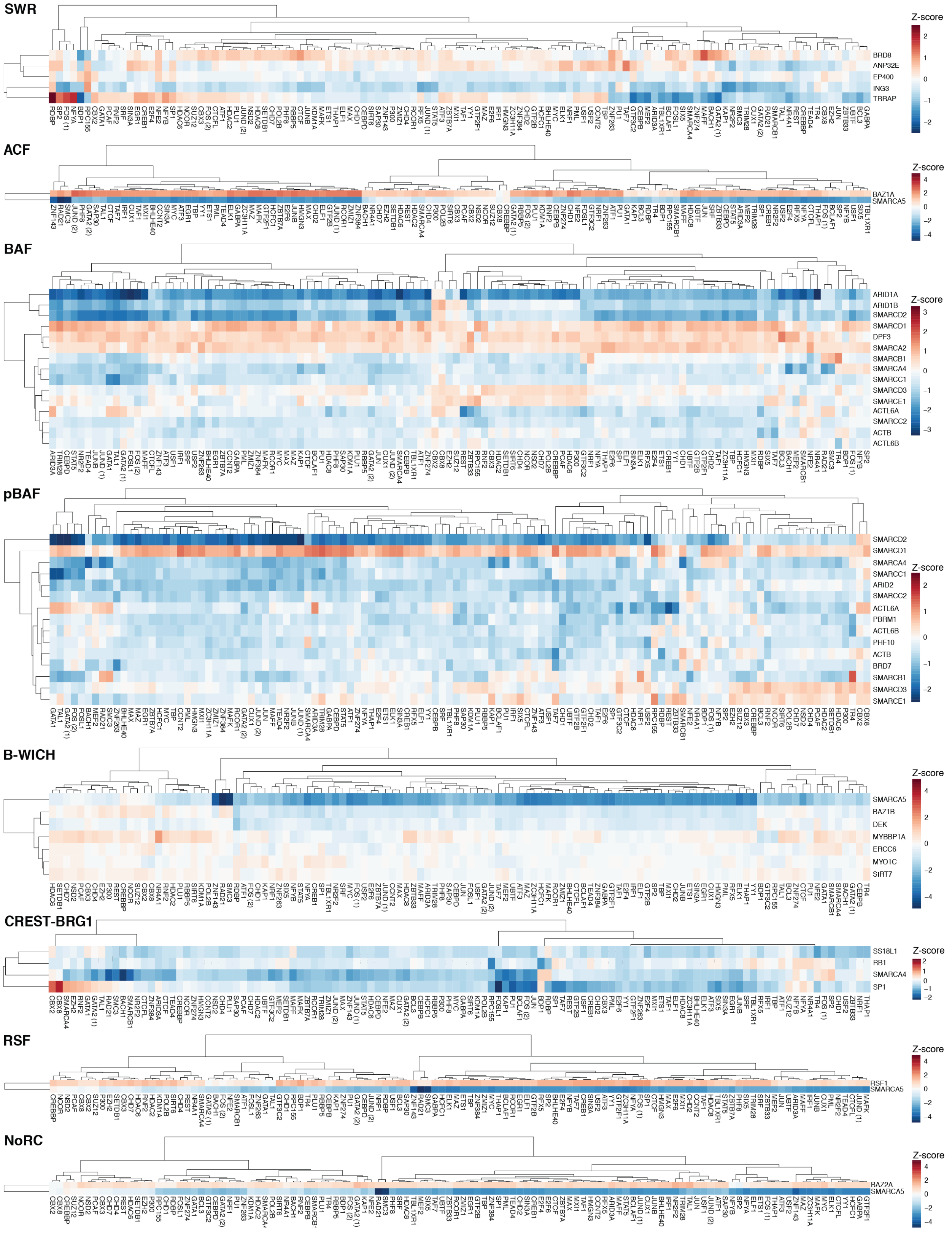


**Supplementary Figure 14. Differential accessibility in TF binding sites.** Heatmaps of chromatin accessibility at transcription factor binding sites (TFBSs) for the different chromatin remodeling complexes targeted. The fraction of reads in TFBSs was calculated for each single cell. Fractions are standardized for each TFBS over all single cells and then averaged over all single cells for each sgRNA cell population. The accessibility *Z*-scores are computed over all cells in the screen. Subunits with less than 50 single cells are not shown. Complexes are ordered by mean accessibility *Z*-score (mean over TFBS) from those with the largest increase in accessibility to those with the largest decrease in accessibility.

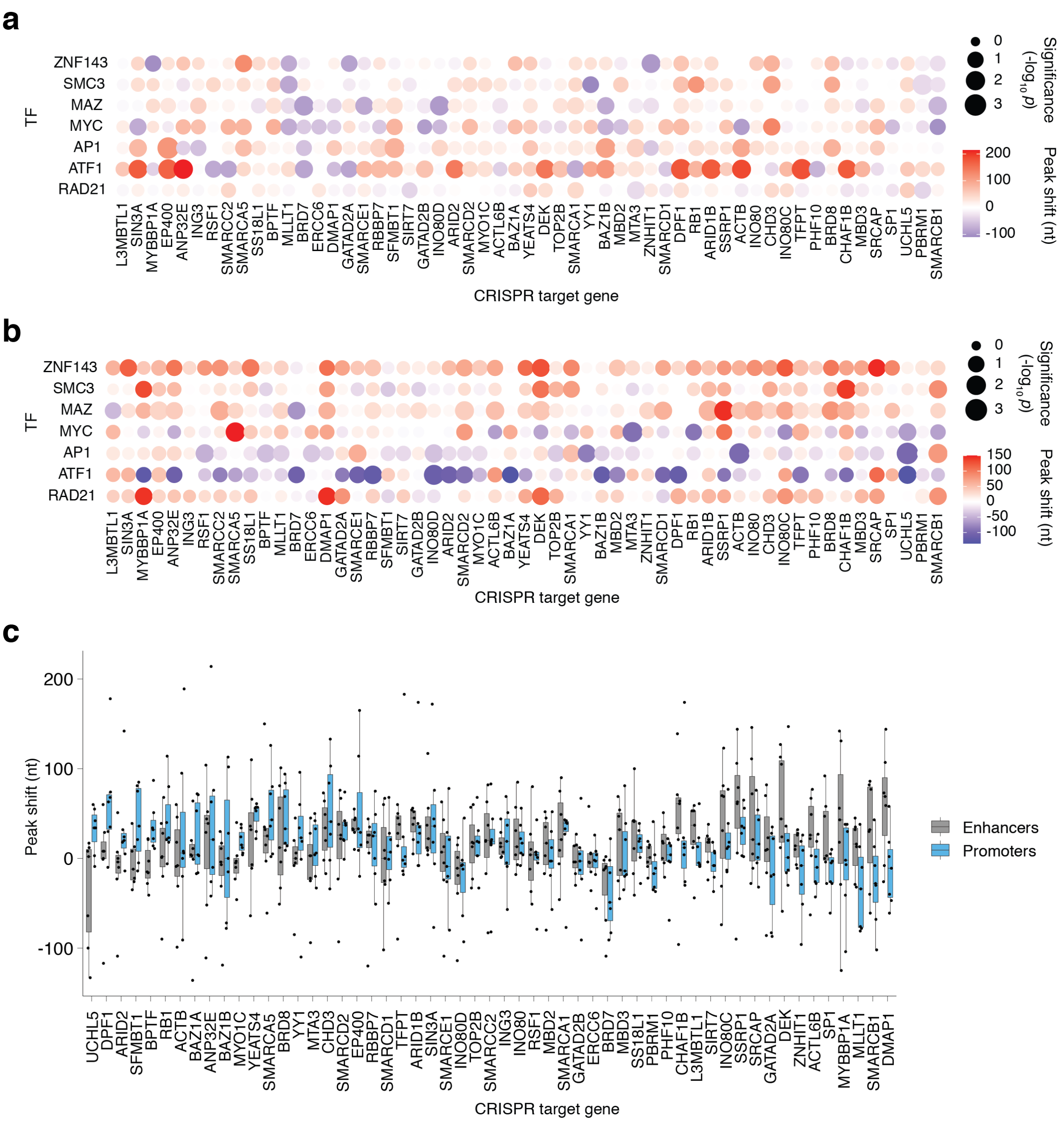


**Supplementary Figure 15. Nucleosome shifts around TFBSs in promoters and enhancers.** **(a, b)** Peak shifts in promoters (**a**) or enhancers (**b**) at the TFBS shown on the *y*-axis in cells that receive a sgRNA targeting the gene shown on *x*-axis. The color of the bubble corresponds to the peak shift (nt) and the size of the bubble represents the empirical *p*-value calculated by a label permutation test. **(c)** Boxplots summarizing the peak shifts in TFBSs located in enhancers and promoters for cells that receive a sgRNA targeting the gene shown on *x*-axis. Each dot represents a different TFBS from the 7 TFBS with symmetric nucleosome positioning[12].

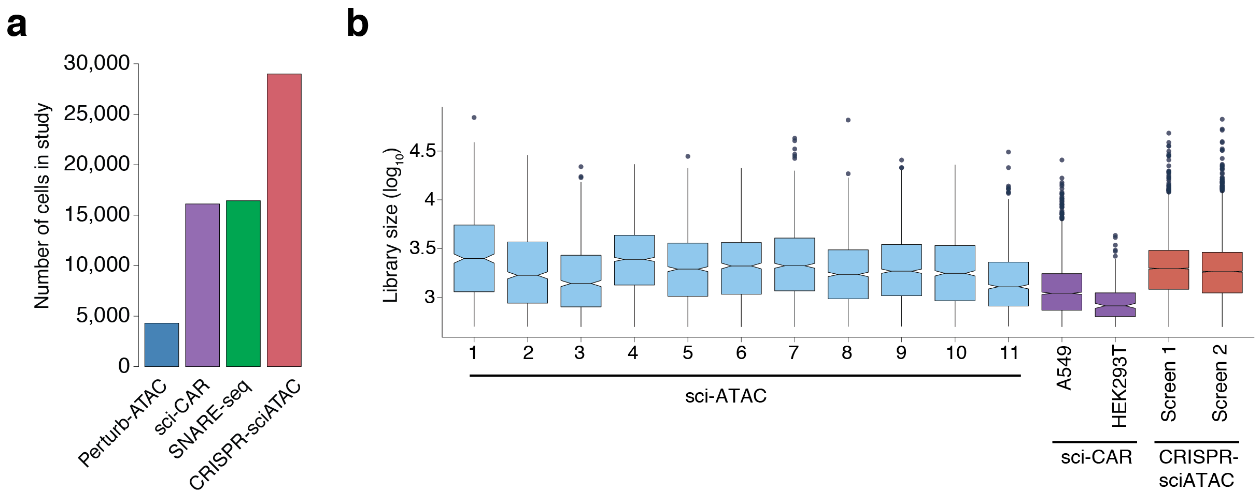


**Supplementary Figure 16. Comparison of CRISPR-sciATAC to prior related studies. (a)** Number of cells profiled in CRISPR-sciATAC and other studies that capture ATAC with a second modality: Perturb-ATAC[13], sci-CAR[14] and SNARE-Seq[15] **(b)** Number of ATAC reads per cell in datasets from published sci-ATAC studies[14,16].

### Supplemental sequences

TnY coding sequence (ViPAR P50K, M53Q) - No STOP codon

```
ATGACCCACTCCGATGCGAAACTGTGGGCTCAGGAGCAATTCGGTCAGGCCCAACTGAAAGATCCGCGCCGCACCCA
GCGCCTGATTTCTCTGGCGACCAGCATTGCTAACCAGCCGGGTGTTAGCGTTGCGAAACTGCCGTTTTCTAAAGCCG
ATCAGGAGGGCGCGTACCGTTTCATTCGTAACGATAACATCGACGCGAAAGACATCGCTGAAGCAGGCTTTCAGTCC
ACCGTATCCCGCGCTAACGAACACAAAGAGCTGCTGGCGCTGGAAGACACTACGACCCTGTCTTTCCCGCATCGTTC
CATCAAAGAAGAACTGGGCCATACGAACCAGGGTGATCGCACCCGCGCCCTGCACGTTCACTCTACCCTGCTGTTCG
CGCCGCAGAACCAGACTATCGTGGGTCTGATCGAGCAGCAGCGTTGGTCTCGTGATATTACTAAACGCGGTCAGAAA
CATCAGCACGCTACCCGTCCTTATAAAGAAAAAGAATCCTATAAATGGGAGCAGGCTTCCCGTCGTGTTGTGGAGCG
CCTGGGTGATAAAATGCTGGATGTCATTTCTGTTTGCGACCGCGAGGCAGATCTGTTTGAATACCTGACCTACAAAC
GTCAACACCAGCAGCGTTTCGTTGTTCGTAGCATGCAGTCTCGCTGTCTGGAAGAACACGCTCAGAAACTGTATGAC
TACGCACAGGCGCTGCCATCTGTAAAAACGAAGGCACTGACCATCCCTCAAAAAGGTGGCCGTAAAGCACGTGACGT
TAAACTGGACGTTAAATACGGCCAGGTTACTCTGAAAGCGCCGGCCAACAAAAAGGAGCACGCAGGCATTCCGGTTT
ACTACGTGGGCTGCCTGGAACAGGGTACTTCCAAAGATAAACTGGCGTGGCACCTGCTGACCTCTGAACCTATTAAC
AACGTCGAGGATGCCATGCGTATCATCGGCTACTACGAACGTCGTTGGCTGATCGAGGATTTTCACAAAGTATGGAA
ATCCGAAGGTACTGACGTAGAATCCCTGCGTCTGCAGAGCAAAGACAACCTGGAACGTCTGTCCGTTATCTACGCGT
TTGTTGCTACCCGCCTGCTGGCACTGCGTTTTATCAAGGAAGTTGATGAACTGACCAAAGAAAGCTGTGAAAAAGTT
CTGGGCCAGAAAGCGTGGAAACTGCTGTGGCTGAAGCTGGAATCTAAAACCCTGCCGAAAGAGGTACCGGACATGGG
TTGGGCTTATAAAAACCTGGCTAAACTGGGTGGCTGGAAGGACACTAAGCGTACCGGTCGCGCTTCTATCAAAGTTC
TGTGGGAGGGTTGGTTCAAACTGCAGACCATCCTGGAGGGCTATGAACTGGCGATGTCCCTGGACCAC
```
